# Virus-host interactions and viral population dynamics across atmospheric cloud events

**DOI:** 10.64898/2026.05.18.725630

**Authors:** Janina Rahlff, Naama Lang-Yona, Ella Lahav, George Westmeijer, Ritam Das, Katrin Buder, Rebecca Bueschel, Julia Micheel, Sabine Eckhardt, Nikolaos Evangeliou, Christine Groot Zwaaftink, Manuela van Pinxteren

## Abstract

**Background:** Cloud water harbors diverse microbial communities despite its extreme oligotrophic conditions. However, the ecological and evolutionary dynamics of viruses in these transient atmospheric habitats remain poorly understood. Clouds have traditionally been regarded primarily as passive carriers of microorganisms rather than as active ecological environments supporting microbial interactions. In this study, cloud water was sampled at Mount Verde, Cape Verde Islands (744 m a.s.l.). We performed metagenomic analyses of iron-flocculated cloud water alongside genome analyses of a bacterial isolate and metagenome-assembled genomes using established bioinformatic approaches. Viral diversity, virus-host interactions, metabolic functions, genetic adaptations, and viral population dynamics across cloud events were investigated. In addition, UV-B resistance experiments were conducted for a novel cloud-water isolate.

**Results:** We isolated 24 cloud water bacteria, including four novel species lineages, and recovered 62 high-quality metagenome-assembled genomes, including 10 novel species lineages. We identified 458 viral operational taxonomic units and 237 virus-host linkages across diverse prokaryotic hosts, revealing active viral predation across diverse bacterial taxa. In addition, CRISPR spacer matches from isolates of novel bacterial lineages such as *Deinococcus nubigenus* MPC36 were found. Viruses carried genes involved in host adaptation to environmental stressors, including cold-shock response, UV radiation resistance, and osmotic stress. In addition, viral populations exhibited SNP-level microdiversity and shifts in single-nucleotide variant composition across temporally proximate cloud events, indicating rapid population turnover. Experimental characterization of the cloud isolate *Curtobacterium nubigenum* MPC39 further revealed pronounced resistance to UV-B radiation and the presence of an inducible prophage, Curtobacterium phage vB_CnuS_Cirrus1 assigned to the new viral family *Nebulaviridae*, which could be validated in transmission electron microscopy. Reconstructed genomes from cloud-associated bacteria encoded carbon monoxide dehydrogenase genes and UV resistance genes, suggesting trace gas metabolism and enhanced UV protection as survival strategies in oligotrophic cloud droplets. In silico replication rates estimated using iRep were consistent with active bacterial replication at the time of sampling.

**Conclusions:** Together, these findings demonstrate that clouds are not merely passive carriers of microorganisms, but dynamic atmospheric ecosystems in which virus-host interactions shape microbial diversity and contribute to microbial turnover, atmospheric dispersal, and cloud-water biogeochemistry.

## Introduction

The atmosphere is estimated to contain 5 × 10^22^ bacterial and archaeal cells [1]. Clouds, as one important atmospheric system, have been recognized as unique habitats or “oases” for microorganisms [2, 3]. As such, they serve as conduits for the dispersal of microbes across diverse ecosystems, impacting biodiversity and ecological dynamics on a global scale [4, 5]. Composed of tiny water droplets or ice crystals suspended in the air, clouds form when moist air rises, cools, and condenses. However, as a microbial habitat, they have to be considered extreme, due to exposure to ultraviolet (UV) radiation, low pH, low nutrients, and cold temperatures [4]. Despite these extreme conditions, microbes remain metabolically active and can thrive [6, 7] while chemical and microphysical factors in clouds may induce oxidative, osmotic, and UV-related stress [8].

This paradox of clouds as both harsh and life-supporting habitats highlights their potential as transient yet functionally relevant microbial ecosystems. Despite the significance of clouds as microbial environments, research on the role of viruses and their prokaryotic hosts in these ecosystems is limited. Only a handful of studies have reported viruses in cloud samples, including plant-infecting viruses such as the Tomato Mosaic Tobamovirus [9], and different DNA and RNA viruses [10], although a recent estimate on the size of the cloud virome was at ∼10^21^ virus particles globally [11]. One major research question is whether the atmosphere is a transport medium or functions as a habitat for an active and permanently resident microbiome [12], i.e., can be considered a true ecosystem. Uncovering predator-prey dynamics between viruses and their microbial hosts could redefine clouds from passive reservoirs into active ecological networks. Critical aspects of viral ecology, such as viral diversity, host interactions, and the overall impact of viruses on airborne microbes, are largely unexplored. The challenge of studying viruses in the atmosphere and clouds is compounded by the ultra-low biomass of these environments [3, 13]. Although amplification-based molecular techniques like 16S rRNA gene sequencing are often the method of choice when faced with low DNA yields, they are not effective for viral analysis [14]. Consequently, our understanding of airborne viruses remains underdeveloped. Recent global events, such as the SARS-CoV-2 pandemic, have shifted attention toward airborne pathogens and their transmission, particularly in indoor environments [15, 16]. This has led to advancements in air sampling technologies and methods aimed at detecting infectious viruses [17]. Within the atmosphere and clouds, viruses could have important functions, for instance, viruses may act as ice-nucleating particles [18, 19], and as such could lower the freezing point of water in the atmosphere. This function may significantly influence cloud formation and precipitation processes, as the presence of these viral particles can enhance ice crystal formation at higher temperatures than would normally be possible [18]. Previous studies suggest that airborne viruses may contribute to the dispersal of auxiliary metabolic genes, including those relevant for resistance [20, 21]. Viruses within clouds must withstand a range of environmental conditions during their residence in cloud droplets, including fluctuations in pH, relative humidity, and temperature, which may affect their viability and activity (reviewed by Longest, et al. [22]). Overall, a deeper investigation into the presence and role of viruses in cloud water is essential for understanding their contributions to atmospheric processes, their potential to disperse into new environments, and their capacity to establish ecological interactions.

Here, we investigated virus-host interactions across successive cloud events by sampling cloud water on Mt. Verde (744 m a.s.l.) in the Cape Verde Islands (Fig. 1). We applied shotgun metagenomics in combination with bacterial isolation and resistance assays. We hypothesized that bacterial viruses (phages) and their interactions with prokaryotic hosts would be detectable, consistent with previous studies reporting their presence in clouds. Given the low abundance of bacterial cells in cloud water, we expected viruses to be primarily associated with the most abundant bacterial species. Furthermore, we anticipated that bacterial and viral communities in cloud water would be characterized by pronounced inter-sample heterogeneity, reflecting the transient nature of this habitat. Finally, we predicted that functional diversity, particularly adaptive genes, would reflect the harsh environmental conditions to which these microbes and viruses are exposed.

**Figure 1:**
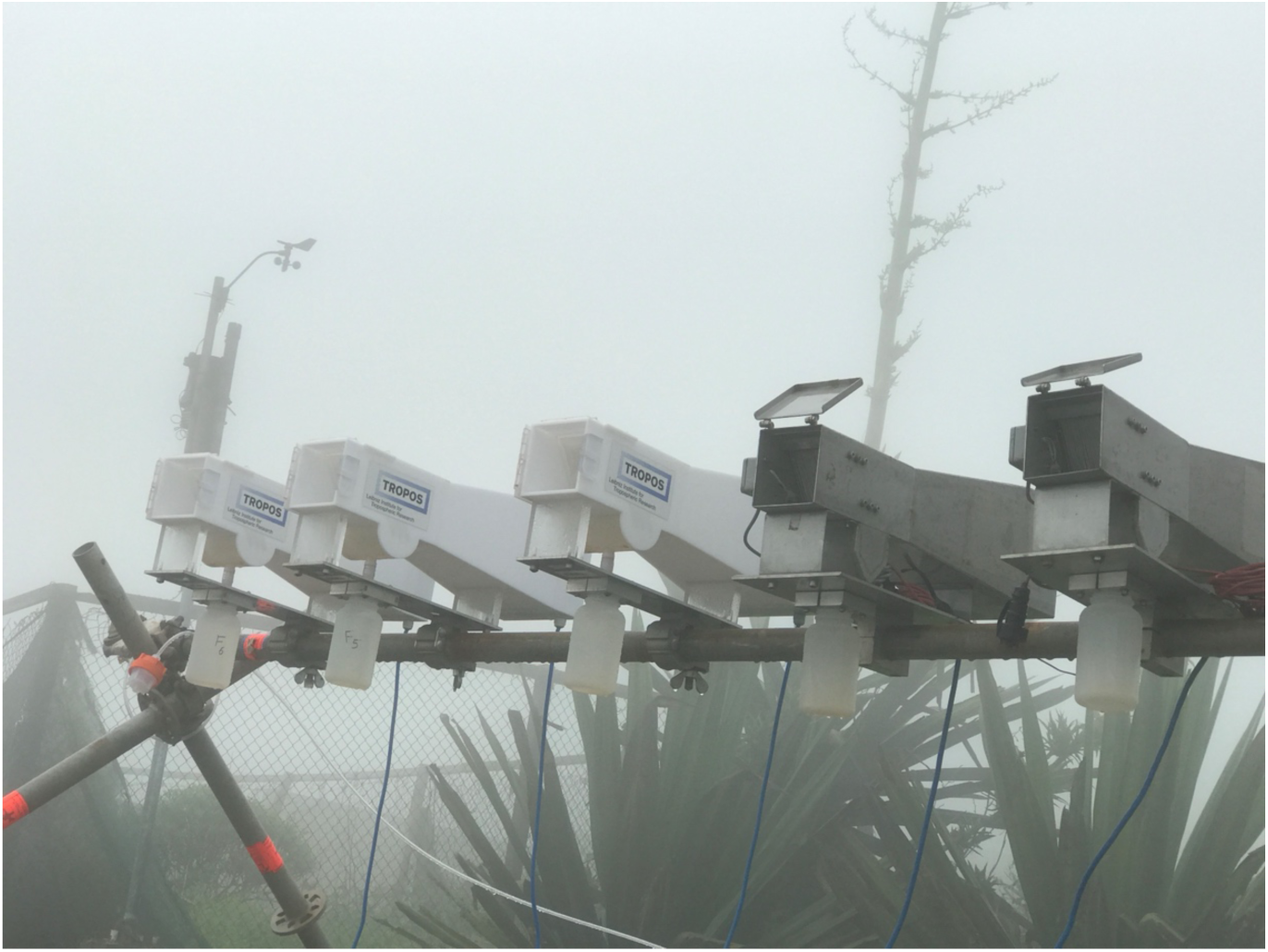
Caltech active strand cloud water collectors during sampling on Mt. Verde. (Photo: René Rabe, TROPOS Leipzig, with permission)

## Methods

### Sampling of cloud water

Cloud water sampling was performed on Mt. Verde on São Vicente Island (16.8695° N, 24.9338° W) during the MarParCloud campaign [23] from 20 September to 5 October 2017, using six compact Caltech active strand cloud water collectors (CASCC2, Fig. 1) [24]. The collectors, comprised of four plastic and two stainless steel units, operated in parallel for sampling durations ranging from 2.5 to 13 h, yielding between 78 and 544 mL of cloud water per sampler. Mt. Verde, 744 m above sea level and located in the northeast of São Vicente Island, was enveloped in clouds approximately 58 % of the campaign duration, with longer cloud coverage observed at night. Aerosol particle samplers ran continuously during cloud events, and the cloud droplets were effectively separated from the aerosol particles, ensuring accurate measurements of both cloud water and aerosol composition. Between samplings, the cloud water samplers were washed with MilliQ, and one such sample was collected as blank (BW13). The six cloud water samples (WW3, WW8, WW16, WW25, WW29, and WW85) and the blank sample (BW13) were stored at -20 °C until from 2017 until 2023.

### Bacterial isolation, plaque assay, DNA extraction, sequencing and genome analysis

Samples were thawed in a water bath at ∼30 °C, and 10 mL of sample was concentrated using centrifugation at 4000 × g for 10 min in an Amicon® Ultra -15 Centrifugal filters Ultracel ® - 10K (Merck Millipore, Burlington, MA, USA). Concentrated samples were plated undiluted and in serial dilutions of factor 10 in Reasoner’s 2A (R2A, Dinkelberg Analytics GmbH, Gablingen, Germany) and Luria*-*Bertani (LB) broth on corresponding agar plates. Emerging bacteria were picked from the plates and purified by replating thrice, stocked for long-term storage at -80 °C and an overnight culture was pelleted for DNA extraction. Bacterial genomic DNA was extracted using the E.Z.N.A. ® Tissue DNA kit (Omega Bio-tek, Norcross, USA) and quantified using the Qubit dsDNA HS Assay Kit on a Qubit 3.0 Fluorometer (Life Technologies / Thermo Fisher Scientific). Concentrated cloud water samples were tested in plaque assays against overnight cultures of the bacterial isolates to find lytic phage plaques (see Supplementary Material). A list of isolates and genome information can be found in Supplementary Table S14.

### Whole-genome sequencing and analysis

Library preparations were performed using the Nextera XT DNA Library Preparation Kit (Illumina, San Diego, CA, USA) with modifications as described in [25]. In brief, DNA was treated with Tn5 transposase. After the Tn5 reaction was stopped, Nextera index primers (Illumina) were added, and the samples were PCR-amplified for 12 cycles. The amplified samples were cleaned using SPRIselect beads (Beckman Coulter, Brea, CA, USA) and subsequently quantified using Qubit dsDNA HS Assay test. Library size distributions were analyzed using the TapeStation D5000 ScreenTape assay (Agilent Technologies, Santa Clara, CA, USA). Short-read sequencing of the DNA from 24 isolated strains was performed on a NovaSeq6000 platform using a NovaSeq 6000 SP 500 cycles v.1.5 flow cell at the inhouse sequencing facility of the FLI. Reads were trimmed using bbduk as part of BBMap v.38.22-0 [26], quality-controlled in fastqc v.0.12.1 [27] and subsequently assembled using metaSPAdes v.3.15.3. [28] using the --isolate option. After a size filtering step (>1000 bp), CheckM2 v.1.0.2 [29] was used for quality check of resulting genomes and the classify_wf of GTDB-Tk v.2.1.1 was applied for taxonomic assignment with r207_v2 database [30]. All non-prokaryotic genomes were removed from further analysis. Prophages were searched using VIBRANT v.1.2.1 [31] and PHASTEST v.1.1 [32] within Proksee v.1.1.0 [33]. Functional annotations were performed using DRAM v.1.5.0 [34]. CRISPRCasFinder v.4.2.20 [35] within Proksee v.1.1.0 [33] was run to detect CRISPR systems, the adaptive immune system of prokaryotes [36], and associated CRISPR spacers from evidence level 4 arrays were extracted.

### Iron flocculation, DNA extraction and library preparation

Assuming overall low biomass in the cloud water samples, the complete remaining sample volume was treated with iron-III-chloride to flocculate viruses and bacteria at a concentration of 10 mg L^-1^ FeCl_3_ × 6H_2_O, which is ten times higher than the concentration used in the standard protocol [37] and that was recommended for freshwater samples [38]. The flocs were then captured using polycarbonate track-etched membranes (0.2 µm pore size, 47 mm diameter, Whatman Nucleopore, Maidstone, UK) combined with a vacuum pump PC 2004 VARIO (VACUUBRAND GmbH, Wertheim AG, Germany). DNA was extracted from these membranes using the DNeasy Power Soil Pro Kit (Qiagen, Hilden, Germany), quantified with a Qubit 3 Fluorometer and further treated with the Genomic DNA Clean & Concentrator®-10 Kit (Zymo Research Europe GmbH, Freiburg im Breisgau, Germany). The sample WW85 was treated slightly different and purified and concentrated with the DNA Clean & Concentrator^TM^-5 Kit (Zymo Research Europe GmbH), which is only suitable for DNA size limits of 50 bp to 23 kb. As for the strains genomes, library preparations were performed using the Nextera XT DNA Library Preparation Kit (Illumina) as explained for the whole-genome sequencing. Short-read sequencing of the metagenomic DNA was performed on a NovaSeq 6000 platform at the inhouse sequencing facility of the Leibniz Institute on Aging - Fritz Lipmann Institute (FLI).

### Viral metagenomic analysis

Paired-end reads were adapter-trimmed as described above and underwent removal of low-quality reads in Sickle v.1.33. The trimmed reads were assembled with metaSPAdes v.3.15.5, MetaviralSPAdes v.3.15.5 [39], and MEGAHIT v.1.2.9 [40]. The generated scaffolds were filtered for small fragments (1000 bp) and combined for each sample. Mobile genetic elements, including viruses, were identified using GeNomad v.1.5.2. [41] with database v.1.3., VIBRANT v.1.2.1 and VirSorter2 v2.2.4 with setting --include-groups “dsDNAphage,ssDNA,NCLDV,lavidaviridae” [42]. All viral scaffolds detected were combined and filtered for size (>10 kb). VIRIDIC v.1.0_r3.6 [43] was run to cluster viral scaffolds with 95 % intergenomic similarity. Only one viral operational taxonomic unit (vOTU) of each species cluster was kept, preferentially a circular scaffold or the longest one. The vOTUs underwent quality checks in CheckV v.1.0.1. with database v.1.5 [44]. Genes were predicted using Prodigal v.2.6.3 in meta mode [45] and GC content using emboss v.6.6.0 with settings infoseq -auto -only -name -length -pgc stdin. Protein-based viral clustering with a reference database (1Mar2024) from INPHARED [46] was conducted using vConTACT2 v.0.11.3 [47]. The results were compiled using graphanalyzer.py v.1.6.0 [48]. Integrated Phage Host Prediction (iPHoP) v.1.4.1 was run against database vJun_2025_pub_rw to match vOTUs to a potential host at the genus level [49]. CRISPR spacers obtained from the bacterial isolates were matched using blastn v2.14.0 --short algorithm with an 80 % similarity filtering step to the dereplicated set of vOTUs and the plasmids previously identified by GeNomad and dereplicated using Cd-hit v.4.8.1 [50] with 99 % identity. Mapping to vOTUs was conducted using Bowtie2 v.2.5.1 [51] with subsequent settings for mapping 90 % identical reads: --ignore-quals --mp 1,1 --np 1 --rdg 0,1 --rfg 0,1 --score-min L,0,-0.1 -1 [52]. The scripts calc_coverage_v3.rb [53] and calcopo.rb [54] were used to calculate read coverage and read breadth, respectively. Breadth was calculated to comply with viromics standards that 75 % of the vOTUs have at least a coverage of >=1 [55]. Coverages were normalized to sequencing depth. Relative abundance represents each normalized coverage value as a percentage of the total. Viral micro-diversity was investigated using profiles from inStrain v.1.9 [56]. Instrain was used in profile mode with default settings on the Bowtie2 mappings (see above), following conversion to bam files with samtools v.1.17 [57], and then in compare mode for comparisons across all samples. The vOTUs were functionally annotated using DRAMv v.1.5.0 [34].

### Taxonomic profiling of the prokaryotic community, binning of MAGs and taxonomic assignment

Read-based taxonomic profiling of the cloud water prokaryotic community was performed using motus v.3.1.0 [58]. Diversity analysis (Shannon-Wiener index, non-metric dimensional scaling) was performed using the package vegan v.2.7.1 [59] within the R programming environment v.4.4.0 [60]. In addition, metagenome-assembled genomes (MAGs) were binned from the unfiltered MEGAHIT and metaSPAdes assemblies with part of the nf-core/mag v.3.2.1 pipeline [61, 62] that uses Nextflow v.24.10.3 [63]. This included the tools MaxBin2.0 [64], MetaBAT 2 [65], and CONCOCT [66] for binning. The following steps were then conducted manually: MAGs were aggregated using DAS tool v.1.1.3 [67] and manually refined in uBin v.0.9.20 [53]. The resulting bacterial MAGs were quality-checked with CheckM2 v.1.0.2 and taxonomically classified with GTDB-Tk v.2.1.1 combined with GTDB release 214. Together with the genomes of the isolates (see above), the MAGs were dereplicated in dRep v.3.5.0. [68]. The trimmed reads were mapped to the dereplicated set of isolate genomes and MAGs using Bowtie2 v.2.5.1. (using --reorder flag). To find evidence for in situ replication of cloud water bacteria, indices of replication were predicted using iRep v.1.1.7 [69] with -mm 3 and otherwise default settings. Both the MAGs and the isolate genomes were functionally annotated with KofamScan v.1.3.0 [70]. KEGG module completeness was assessed with the Reconstruct tool of the KEGG mapper (v. July 2024). These annotations were used for detection of metabolic pathways, genes for the subunits of aerobic carbon monoxide dehydrogenase (CODH), UV resistance, osmoregulatory, and cold-shock genes. To assess whether MAGs and isolate genomes represent novel species, we first performed a tetra correlation search to identify related reference genomes, followed by ANIb calculation using the JSpeciesWS webserver [71]. A genome was considered a candidate new species if its ANIb to the closest known representative was < 95 %, or if the ANIb was > 95 % but the reference lacked a formal species name, and the species was not assigned by GTDB-Tk classify_wf. MAGs with completeness < 89 % or contamination > 5 % were not proposed as new species, even if the lineage is currently unclassified. New species names have been endorsed by the SeqCode [72].

### Prophage induction

To investigate prophage activation, an induction experiment was carried out for 15 of the isolated bacterial strains. The strains were chosen if they had prophages inserted in their genomes based on analysis with VIBRANT. Cultures were grown overnight and distributed into 24-well plates (Sarstedt, Nümbrecht, Germany) with 500 µL culture per well, and mitomycin C ready-made solution in DMSO (Sigma-Aldrich/Merck, Darmstadt, Germany) was added as a prophage-inducing agent in triplicate at final concentrations of 1, 0.5, and 0.1 µg mL^-1^, alongside untreated controls. Optical density at 600 nm (OD_600_) was monitored hourly using a microplate reader Infinite M1000 Pro (Tecan Group Ltd, Männedorf, Switzerland) to track growth dynamics. A noticeable decline in OD_600_ relative to controls after ∼4 h was considered an indication for successful prophage induction, based on previous experience with psychrophilic bacteria from marine systems [73]. At the conclusion of the assay, supernatants from wells showing lytic activity were harvested, combined, and filtered through 0.2-µm syringe filters. These filtrates were stored at 4 °C for transmission electron microscopy (TEM) analysis and DNA extractions. For the latter, 1 mL of the concentrated sample was subjected to DNA extraction using Wizard PCR DNA Purification Resin and Minicolumns (Promega, Madison, WI, USA) as described previously [74]. The DNA was sequenced using Illumina DNA PCR-free method at SciLifeLab (Solna, Sweden). The sample was sequenced on NovaSeqXPlus (NovaSeqXSeries Control Software 1.3.1.59007) with a 151nt(Read1)-10nt(Index1)-10nt(Index2)-151nt(Read2) setup using ‘25B’ mode flowcell. Trimming was performed as for the other samples, and assembly performed using MetaSPAdes v.3.15.3. and viral scaffolds detected using CheckV v.1.0.1. The resulting viral genomes and the two prophage genomes of the MPC39 strain were investigated in a proteome-based phylogenetic tree, which was built using ViPTree v.4.0 [75] to reveal the induced prophage’s identity. The genome of the phage was visualized in Proksee and compared with phylogenetic neighbors using VIRIDIC [43] and Virclust [76] webtools. In another experiment, prophage induction following UV exposure was assessed throughout 12 h UV exposure intervals (302 nm) per day over a total of 48 h. Induction was evaluated by monitoring culture growth through OD_600_ (Synergy H1 microplate reader), compared to non-exposed cultures (six replica per treatment). Sampling 0.5 mL from each replica in duplicates, OD_600_ was recorded every 12 h throughout the exposure period, and at hourly intervals post-exposure for up to 48 h (Supplementary Fig. S11).

### Transmission electron microscopy analysis

Negative staining followed by TEM was used to visualize the structure of the viruses [77]. In brief, 200 mesh copper, formvar/carbon film coated TEM grids (Science Services GmbH, München, Germany) were glow discharged with a sputter coater Q150T ES (Quorum Technologies, Laughton, East Sussex, UK) to enhance their hydrophilicity. A 10 µL droplet of the virus supernatant after prophage induction was applied onto a sheet of laboratory film (Carl Roth, Karlsruhe, Germany) and a grid with the film layer was placed on. After 1 min of sample adsorption, the liquid was pulled off with filter paper by touching the edge of the grid. Immediately, the grid was put on a droplet of 2 % uranyl acetate solution pH 4 - 4.5 (SERVA Electrophoresis GmbH, Heidelberg, Germany) for 30 s. Excess staining solution was removed with filter paper, and the grid was allowed to dry at room temperature in the dark. Samples were imaged at an acceleration voltage of 80 kV in a JEM-1400 transmission electron microscope (JEOL, Tokio, Japan) connected to a Orius SC 1000A CCD camera (Gatan Inc., Pleasanton, CA, USA) while using the software Gatan Microscopy Suite v.2.31.734.0. Phage head and tail dimensions were measured in ImageJ v.1.54g [78] according to the protocol by Brum [79].

### UV exposure of Curtobacterium nubigenum MPC39

To assess UV radiation effect on *C. nubigenum* MPC39 growth, cells were exposed to UV-B light (302 nm) for 12 h exposure intervals per day over a total duration of four days (and a maximum of 48 h UV exposure), using six 5 mL LB broth replicates at 20 °C. The exposed cultures were then placed on glass slides and stained with SYTO 9 and propidium iodide (LIVE/DEAD BacLight kit, Thermo Fischer Scientific) to visualize live and dead cells, respectively. For each treatment, 4-5 microscopic fields were recorded at 63 × magnification using a Stellaris confocal microscope (Leica, Wetzlar, Germany), and cell counts were analyzed using ImageJ v.1.54s. Post-UV exposure growth curves were monitored by measuring OD_600_ (BioTek Synergy H1 microplate reader, Agilent Technologies, Santa Clara, CA, USA), using a 1:10 dilution of exposed cultures, compared to non-exposed controls (six replicates per treatment). Additionally, colony diameters on LB agar plates, were evaluated following incubation under the same conditions as the liquid cultures exposed to UV-B using a Stemi 508 stereomicroscope (Carl Zeiss AG, Jena, Germany). Images of the biofilm were made with an upright Zeiss Axioscope 5.0.

### Statistical analysis

Significant differences between treatments of UV exposure were tested in R v.2025.09.2 using One Way Analysis of Variance (ANOVA) with the Tukey HSD *post hoc* test with significance set at *p* < 0.01. GraphPad Prism v.10.5.0 was used for statistical analysis of UV and CSP-gene comparisons. CSP and UV-resistance gene data were tested for normality using the Shapiro–Wilk and Kolmogorov-Smirnov tests, and homogeneity of variances was assessed using the Brown-Forsythe test and Bartlett’s test in GraphPad Prism v.10.6.1. The Pearson correlation coefficient was applied to evaluate the relationship between the number of CSP-encoding genes and genome size. Gene density was calculated by dividing the absolute gene number from each genome by their respective genome size. Depending on the results of the normality tests, either one-way ANOVA followed by Tukey’s HSD *post hoc* test or the Kruskal-Wallis test followed by Dunn’s multiple comparisons test was used to compare *uvr* gene number and density among different phyla and across different *uvr* types.

### Backward trajectory analysis for air masses

We conducted 30-day backward trajectory simulations for each cloud-water sample using the Lagrangian particle dispersion model FLEXPART v.10.4 [80]. FLEXPART is an atmospheric transport model designed to simulate the pathways of atmospheric tracers by accounting for advection, turbulent diffusion, convection, and dry and wet deposition processes. It is widely used in the atmospheric sciences for source-receptor analyses, inverse modelling, and the interpretation of in situ and remote-sensing observations. The simulations were driven by operational meteorological fields from the ERA5 reanalysis [81] (hourly, 0.5° horizontal resolution) and initialized with 40000 computational particles released at the sampling location for the respective sampling times. The model tracks the trajectories of these particles backward in time, allowing us to compute emission sensitivity or “footprint” maps that quantify the particle residence time within the planetary boundary layer. These sensitivity fields identify potential source regions and highlight where air masses sampled as cloud water at the site had the strongest surface contact. To characterize the influence of different surface types, we further calculated residence times over specific land-use categories, including ocean, desert, forest, and anthropogenic areas. This separation allowed us to assess the relative contributions of natural versus anthropogenic surfaces and to identify signatures of marine, desert-dust, continental, or pollution-related influence. For the estimation of the dust contribution, the FLEXDUST framework [82] has been applied. Based on soil properties and meteorological conditions dust is suspended and transported. Together, the FLEXPART analyses provide a physically consistent basis for interpreting the chemical composition of the cloud-water samples in terms of their atmospheric transport history and likely source environments.

## Results

### Bacterial and viral communities differ markedly across cloud water samples

We characterized the viral community across all samples to assess diversity, abundance patterns, and taxonomic composition. In total, 458 vOTUs >10 kb length were detected across all samples, of which 263, 58, and 26 were classified as low, medium and high quality (remaining ones not determined), respectively. In total, 49 were proviruses. The guanine-cytosine (G+C) base content of the vOTUs ranged between 27.3 and 74.43 with a median of 64.3. Mapping of reads to the vOTUs revealed that the number of dsDNA phages decreased in the beginning (WW3 & WW8) and then increased again at the end (WW85) of the sampling period. Virophages (Lavidaviridae) were only detected in the last four samples, while the number of giant viruses (nucleocytoplasmic large DNA viruses (NCLDVs) or Nucleocytoviricota) increased during the second half of the campaign (Supplementary Fig. S1). A higher proportion of dsDNA phages was associated with a higher G+C base content of the viral community in each sample (Pearson’s *r* = 0.82, *p* = 0.023, *n* = 7, Supplementary Fig. S1).

The most abundant viral clusters (VC) were VC_1704_0 (27.7 % relative abundance in sample WW25, 14.4 % in sample WW16, and 13.2 % in WW29) and VC_1706_0 (17.3 % in sample WW16, 3.2 % in WW25, Fig. 2a). VC_1704_0 and VC_1706_0 contained vOTUs assigned to NCLDV. VC_1598_0 had 9.9 % relative abundance in WW3, and vOTUs therein showed protein similarities with Microbacterium phage Footloose (Genbank, <u>MZ150789</u>). One vOTU from this VC had the predicted host genus *Rhodococcus* (Actinomycetota). This VC was absent in other samples. VC_786_0 occurred in WW3 and WW8 with 11.3 % and 7.4 % relative abundance respectively, and vOTUs assigned to this cluster had protein similarities to Curtobacterium phage Reje (Genbank, <u>MZ333136</u>) and Actinomyces phage Av-1 (Genbank, <u>DQ123818</u>). A vOTU from this VC was assigned to a *Mycobacterium* sp. host. Sample WW85 contained VC_774_0 with a relative abundance of 11.1 %, which contained two vOTUs with protein similarities to Klebsiella phage ST11-VIM1phi8.3 (Genbank, <u>MK416019</u>). This VC was absent in other samples. The predicted host genera for a vOTU in the cluster were *Pararhizobium* sp. or *Rhizobium* sp. All samples contained singletons and outliers as defined by vConTACT2, which can represent previously unknown viruses. Shannon-Wiener (SW) indices were comparably high in five of the samples with values between 3.5 to 3.7 and at 2.7 for WW16 and 2.9 for BW13 (Fig. 2b). Non-metric multidimensional scaling (NMDS) revealed no clear clustering demonstrating a dynamic viral community (Fig. 2c). Overall, these results highlight a highly dynamic viral community with shifts in dominant viral groups over time and the presence of both well-characterized and potentially novel viruses.

**Figure 2:**
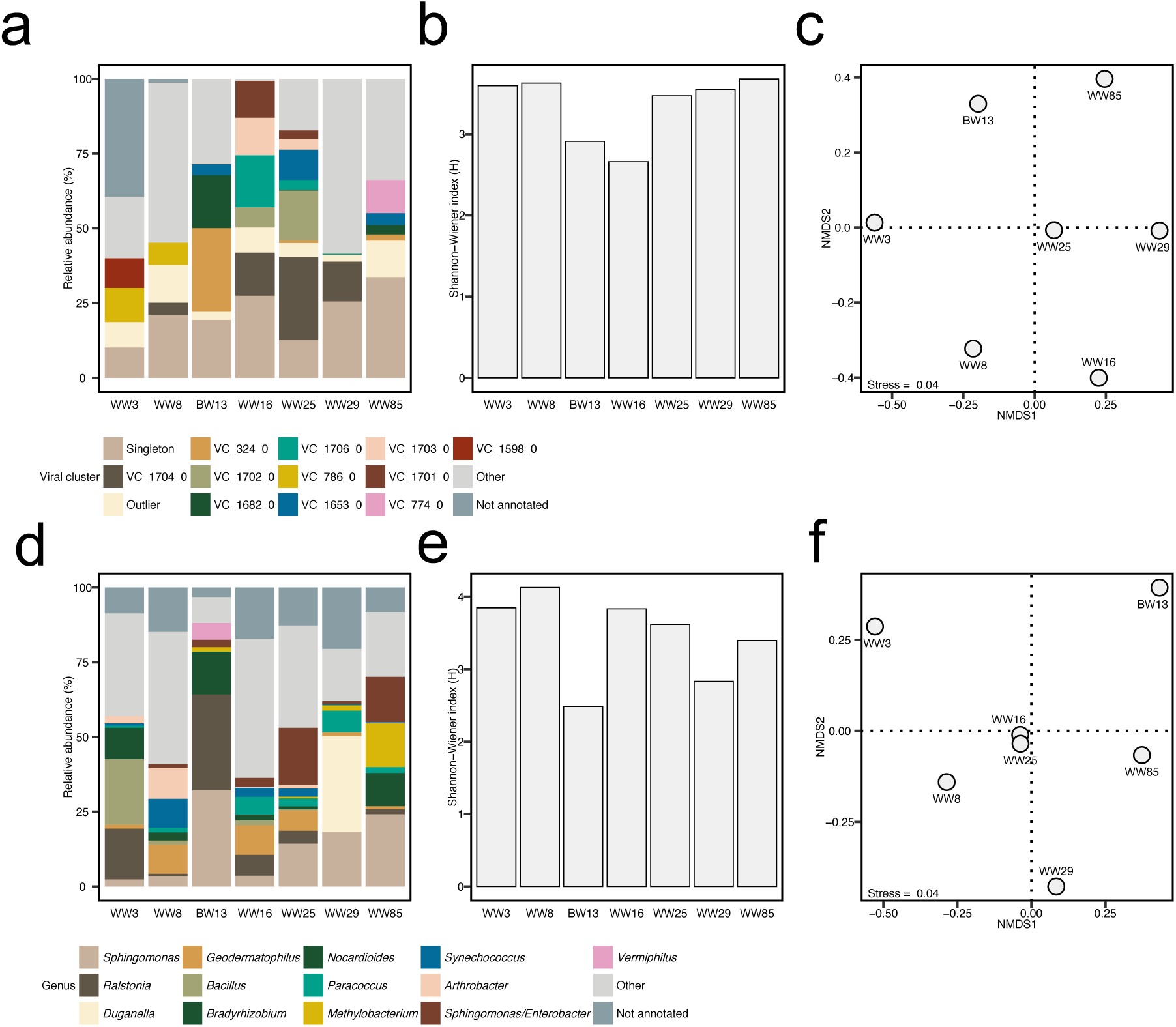
Diversity and composition of the viral and prokaryotic communities. a, Viral community structure, including the 13 most abundant viral clusters while grouping low-abundant taxa as ‘Other’. **b,** Alpha diversity according to the Shannon-Wiener index. **c,** Beta diversity according to non-metric multidimensional scaling (NMDS), based on the relative abundance of the viral OTUs. **d,** Prokaryotic community structure, including the 13 most abundant genera while grouping low-abundant lineages as ‘Other’. **e,** Alpha diversity according to the Shannon-Wiener index. **f,** NMDS based on the relative abundance of the prokaryotic OTUs.

In general, the prokaryotic community was dominated by the bacterial phylum Pseudomonadota (> 50 % of the read counts) with the typically heterotrophic genera *Bacillus, Sphingomonas*, *Ralstonia*, and *Duganella* as the main representatives (Supplementary Table S1). The most abundant autotrophic lineage was the cyanobacterium *Synechococcus* sp., which was detected in 6 of 7 samples. Alpha diversity was similar among the samples with a Shannon-Wiener diversity index between 2.5 and 4.1 (Fig. 2e). Similar to the viral community, the bacterial communities were variable between samples (except for WW16 and 25), as reflected in the community composition (Fig. 2d) and the beta diversity analysis (Fig. 2f). Overall, the prokaryotic communities were dominated by typically heterotrophic bacteria that varied between sampling sites.

### Viruses interact with prokaryotic hosts and evolve in cloud water

Across all samples, 237 virus-host interactions were predicted using iPhoP (Fig. 3). Most vOTUs were assigned to host orders Sphingomonadales (85), Actinomycetales (28), Rhizobiales (24), Cytophagale*s* (19), Burkholderiales (18), Deinococcales (17) and Enterobacterales (14) (Supplementary Table S2). Screening for clustered-regularly interspaced short palindromic repeats (CRISPR) systems in the genomes of the bacterial isolates revealed that *Deinococcus nubigenus* MPC36 and *Sphingomonas panni* MPC37 contained CRISPR arrays. One CRISPR spacer from each strain matched two vOTUs assembled from cloud water sample WW29, which had a high coverage depth (45 - 54, Supplementary Table S3 & S4). Based on read mapping, these vOTUs solely occurred in sample WW29. Two of the targets of the *S. panni* MPC37 CRISPR spacers were identified as prophages. None of the viruses targeted by CRISPR spacers exhibited detectable variants in the micro-diversity analysis. Here, multiple subclusters among the 39 viral strain variant clusters were identified, with variants of eight vOTUs occurring in all samples (Fig. 4a, Supplementary Table S5). Tracking the progression of vOTU clusters across cloud events sampled over time revealed a set of viral variants associated with the earliest sampled events, WW3 and WW8 (20 - 26 September). Investigating the residence time and source region of the air masses for these cloud events, revealed that they had longer residence time over the desert (3.5 vs 2.5 h) and less over the ocean (7 vs 9h) compared to the last two events from WW29 to WW85 (29 September - 5 October, Fig. 4b, Supplementary Fig. S2-S7). Comparison of air mass anomalies (event periods relative to all samples (Supplementary Fig. S2-S7)) highlights distinct transport pathways. During the first episode, air masses primarily originated from Spain, whereas during the second episode, the dominant contribution was from Africa (Supplementary Fig. S8). The WW3/WW8 variant set (26 variant pairs in total) was associated with a major dust event that occurred between 17 and 21 September 2017 (Supplementary Fig. S9 & S10), measured from aerosol samples with 11.6 - 38.2 µg m^-3^ [23]. In contrast, WW29 and WW85 had particulate amounts of 17.2 to 27.3 µg m^-3^ measured from aerosols [23], and these cloud events were associated with longer residence time over the ocean (Fig. 4). The WW3/WW8 variant clusters contained vOTUs linked to host genera *Agromyces* sp. and *Micromonospora* sp. (Actinomycetota) among several unknown hosts. In contrast, vOTUs assigned to *Ralstonia* sp. and *Escherichia* sp. were detected across all samples, while several vOTUs from WW29/WW85 variants had unknown hosts. Air mass trajectory patterns and the presence of dust may explain the distinct viral variant observed at the beginning of the sampling campaign. Overall, virus variant overlap was greater in temporally proximate cloud events, particularly between the first two events.

**Figure 3:**
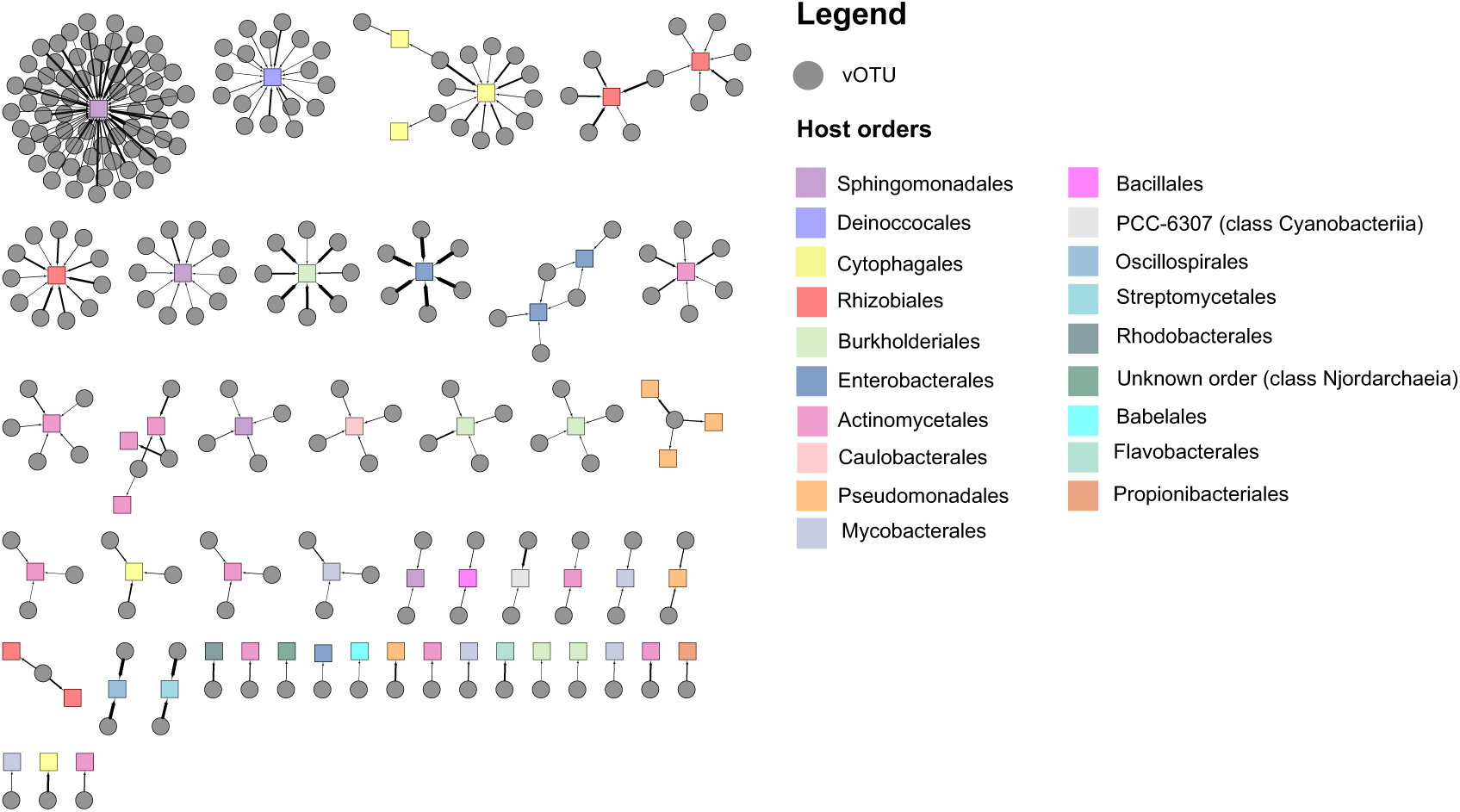
Virus-host interaction network based on iPHoP predictions. Colored squares represent host orders, while black circle represent viral operational taxonomic units (vOTU).

**Figure 4:**
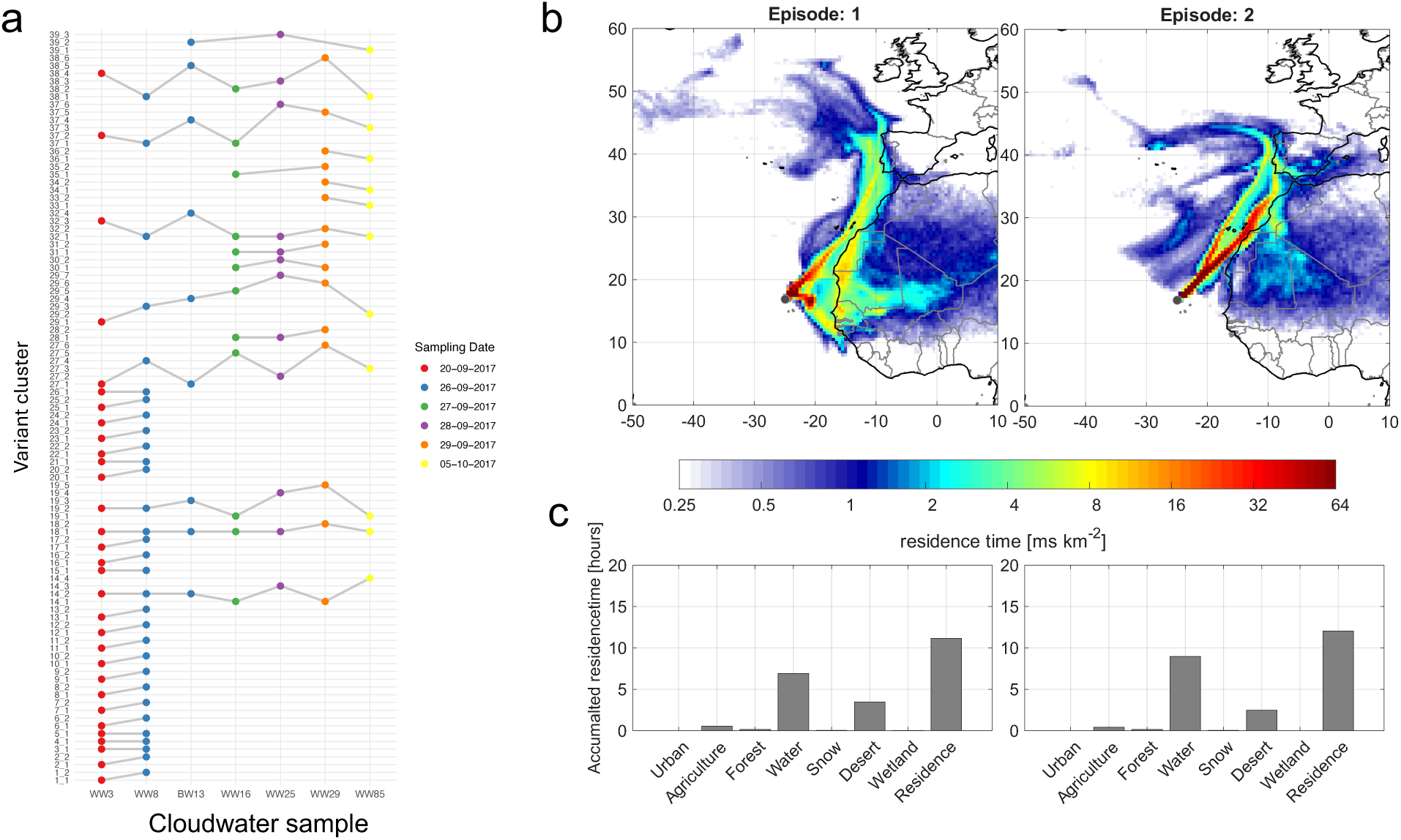
Viral strain-level cluster dynamics across cloud water samples and in relation to air mass source regions. a, The plot shows the occurrence and trajectories of individual vOTUs across multiple samples. Each point represents the cluster assignment of a genome in a specific sample, based on inStrain subcluster analysis. Lines connect the same vOTUs across consecutive samples, allowing visualization of whether a vOTU remains in the same cluster (horizontal line) or is reassigned to a different subcluster (sloped line). The y-axis corresponds to the clusters (1 - 39), while the x-axis represents the samples in chronological order. Points are colored by the sampling date to indicate temporal progression. **b,** These two panels show the average source region (backward trajectories) for episode 1 (20.9.2017, 26.9. 09:00) and episode 2 (28.9.20:00, 5.10.18:40). **c,** The bar plot shows the distribution of emission sensitivities related to different land use categories.

### Functional potential of cloud water prokaryotes and their viruses

The metagenomes yielded 22 de-replicated metagenome-assembled genomes (MAGs). Together with the 16 isolate genomes, this resulted in 38 reconstructed genomes in total, of which 31 had an estimated completeness > 90 % (Supplementary Table 6). These genomes were affiliated with the Pseudomonadota (*n* = 18), Actinomycetota (11), Bacillota (5), Bacteroidota (2), and the Deinococcota (2). Based on MAGs and bacterial isolate genomes, we described several new (Candidatus) species obtained from the cloud water samples (Supplementary Material, Supplementary Table 7). Analysis of the metabolic potential indicated that most lineages had a heterotrophic lifestyle, as only 2/38 genomes encoded for a complete carbon fixation pathway (Fig. 5a). Most taxa obtained their energy by the oxidation of organic compounds, with a widespread genomic potential for the oxidation of organic carbon (e.g. glycolysis and the TCA cycle). In addition, 8/38 genomes encoded for aerobic carbon monoxide (CO) dehydrogenase that would allow these lineages to use the oxidation of atmospheric CO to meet their energy demands in this oligotrophic environment (Fig. 5b).

**Figure 5:**
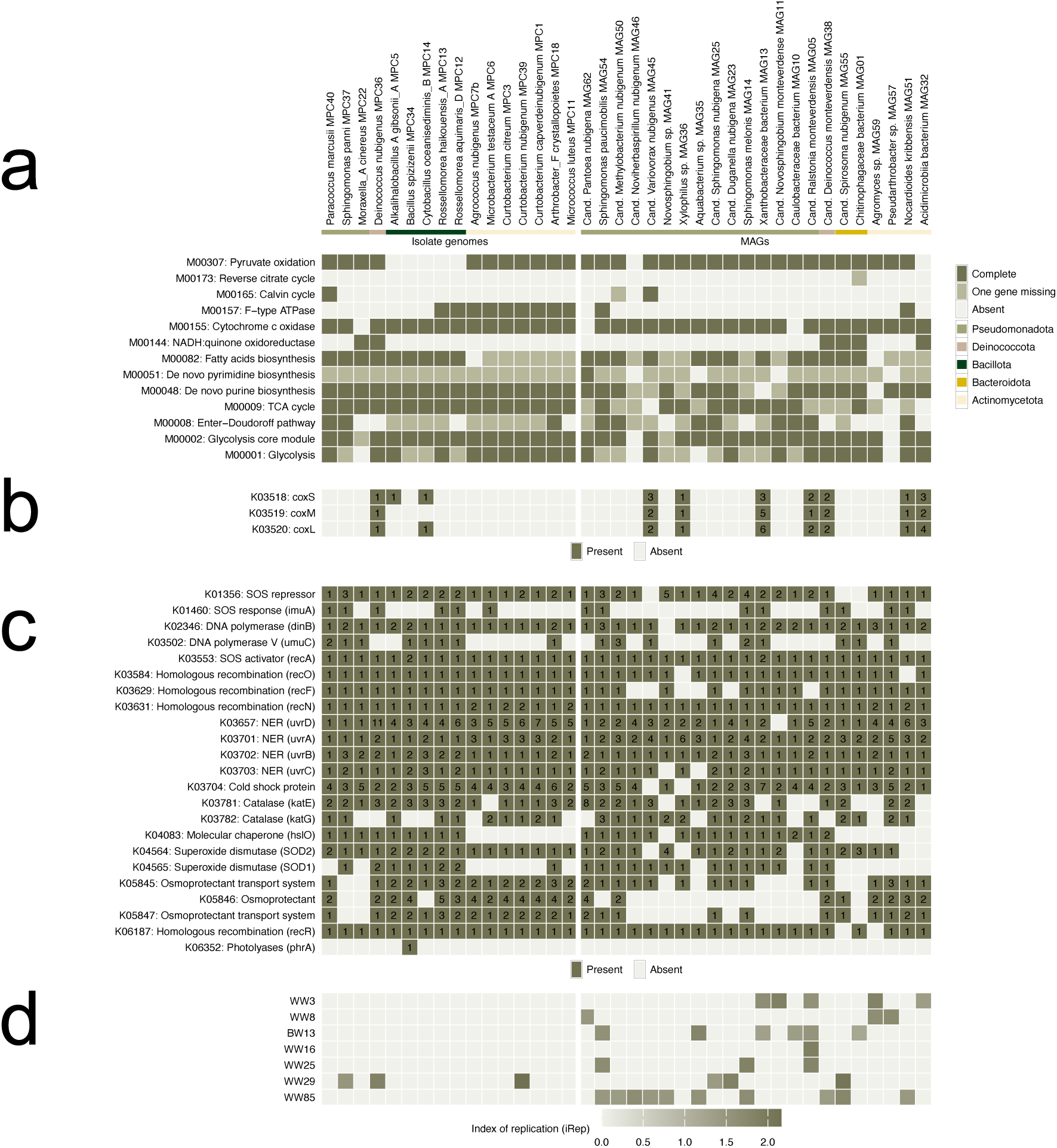
Metabolic potential, adaptive traits and index of replication (iRep) of the de-replicated genomes (isolates and MAGs). **a,** the color code depicts the completeness of the KEGG modules, categorized as complete, one gene missing, and incomplete, referring to KEGG Orthologs (KO) comprising the KEGG modules. **b,** Presence/absence of KEGG orthologs related to the oxidation of carbon monoxide. The number in the colored tiles depicts the copy number of the respective gene in the metagenome assembled genome (MAG). **c,** Presence/absence of Kos involved in various environmental stress related processes, including osmoregulation, oxidative stress, and DNA repair. The number in the tiles depicts the copy number of the respective gene. **d,** Index of replication (iRep) for the de-replicated genomes in each metagenome.

Across several cloud water MAGs and bacterial isolate genomes (Fig. 6), the presence of cold-shock protein (*CSP*)-encoding genes was observed (Fig. 5c). Core DNA repair and SOS response genes (e.g., *recA, recO/F/N, uvrABCD, dinB*) were broadly conserved across the genomes, while stress response elements such as osmoprotectant transporters, catalases, cold shock proteins, and photolyases displayed greater variability in number among strains. The genomes, on average, encoded for approximately 3.18 (± 1.66) *CSPs*, with *Sphingomonas melonis* MAG14 having the greatest number of genes (seven) (Fig. 5c). A significant positive correlation (Pearson’s *r* = 0.36, *p* = 0.026) between the number of *CSP*-encoding genes and the genome size of the bacteria was observed (Fig. 7). Among the genes involved in UV resistance, there were significant differences between the phyla and gene type (Kruskal-Wallis test *p* < 0.0001). Actinomycetota encoded a significantly higher number of *uvrD* genes (4.82 ± 1.25) as compared to *uvrB* (1.82 ± 0.40) and *uvrC* genes (1.10 ± 0.30; Dunn’s multiple comparisons test, *p* < 0.001) and compared to *uvrD* in Pseudomonadota (1.94 ± 1.31) (Dunn’s test, *p* = 0.0053) (Fig. 8a, Supplementary Table 8). Across all genomes, significant differences for the *uvrABCD* gene complex between phyla (ANOVA, F (2, 31) = 8.28, *p* = 0.0013), namely between Pseudomonadota and Actinomycetota (Tukey HSD, *p* = 0.0018) and between Pseudomonadota and Bacillota (Tukey HSD, *p* = 0.049) were observed (Fig. 8b, Supplementary Table 8). These differences were maintained when considering *uvrABCD* gene density, i.e. number of genes per Mb of the genome (Fig. 8c, Supplementary Table 8) across phyla (ANOVA, F (2, 31) = 25.8, *p* < 0.0001), with Actinomycetota having a significantly higher gene density of (2.95 ± 0.56) compared to Bacillota (1.96 ± 0.24, Tukey HSD, *p =* 0.0031), and compared to Pseudomonadota (1.55 ± 0.53, Tukey HSD, *p* < 0.0001). Significant differences between the four gene types across all phyla were also found (Kruskal-Wallis test, *p* < 0.0001). Here, the *uvrD* gene (3.26 ± 2.23) was significantly more abundant than *uvrB* (1.42 ± 0.64) and *uvrC* (1.18 ± 0.56, Dunn’s test, *p* < 0.0001) (Fig. 8d, Supplementary Table 8).

**Figure 6:**
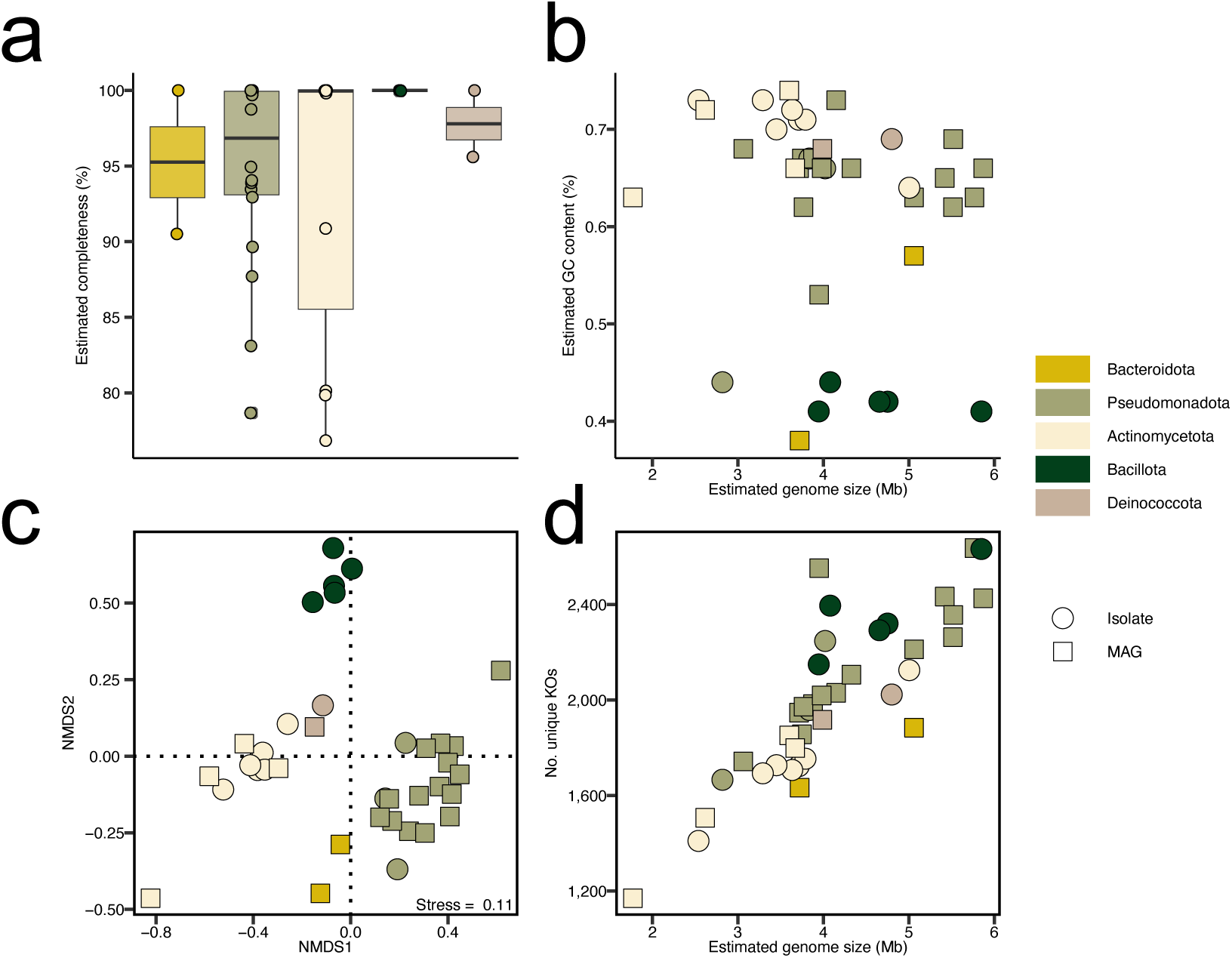
Features of the cloud water-derived metagenome-assembled genomes (MAGs) and genomes of bacterial isolates. Shown are estimated **a,** estimated completeness, **b,** estimated G+C content (%), **c,** grouping of genomes using non-metric dimensional scaling (NMDS) on the repertoire of KEGG orthologs in the individual genomes, and **d,** number of unique KEGG orthologs versus genome size.

**Figure 7:**
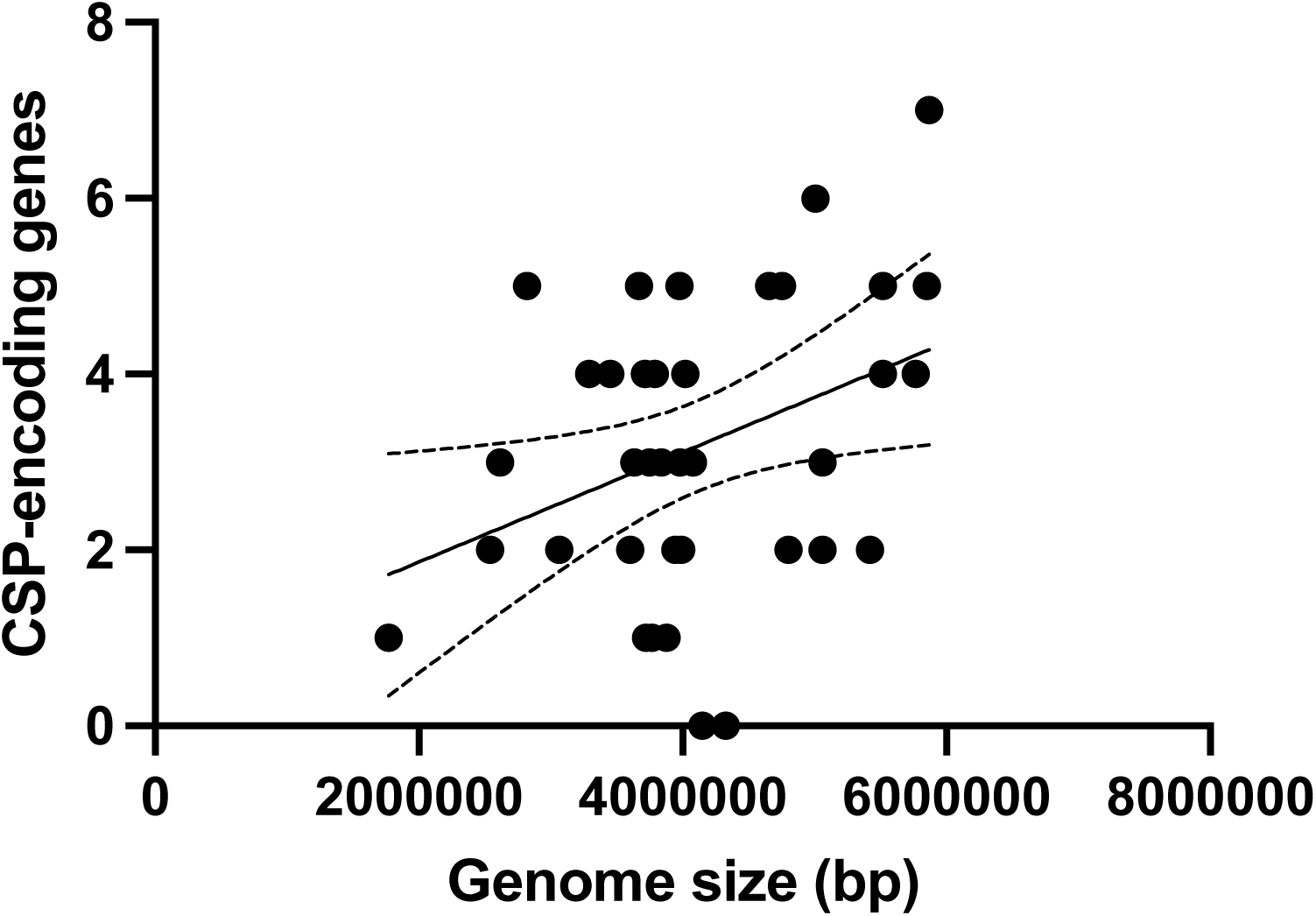
**Correlation analysis of CSP-encoding genes and genome size**. The total number of CSP-encoding genes was found to correlate positively with the genome size of the bacterial strains Pearson’s *r* = 0.36, *p* = 0.026, *n* = 38).

**Figure 8:**
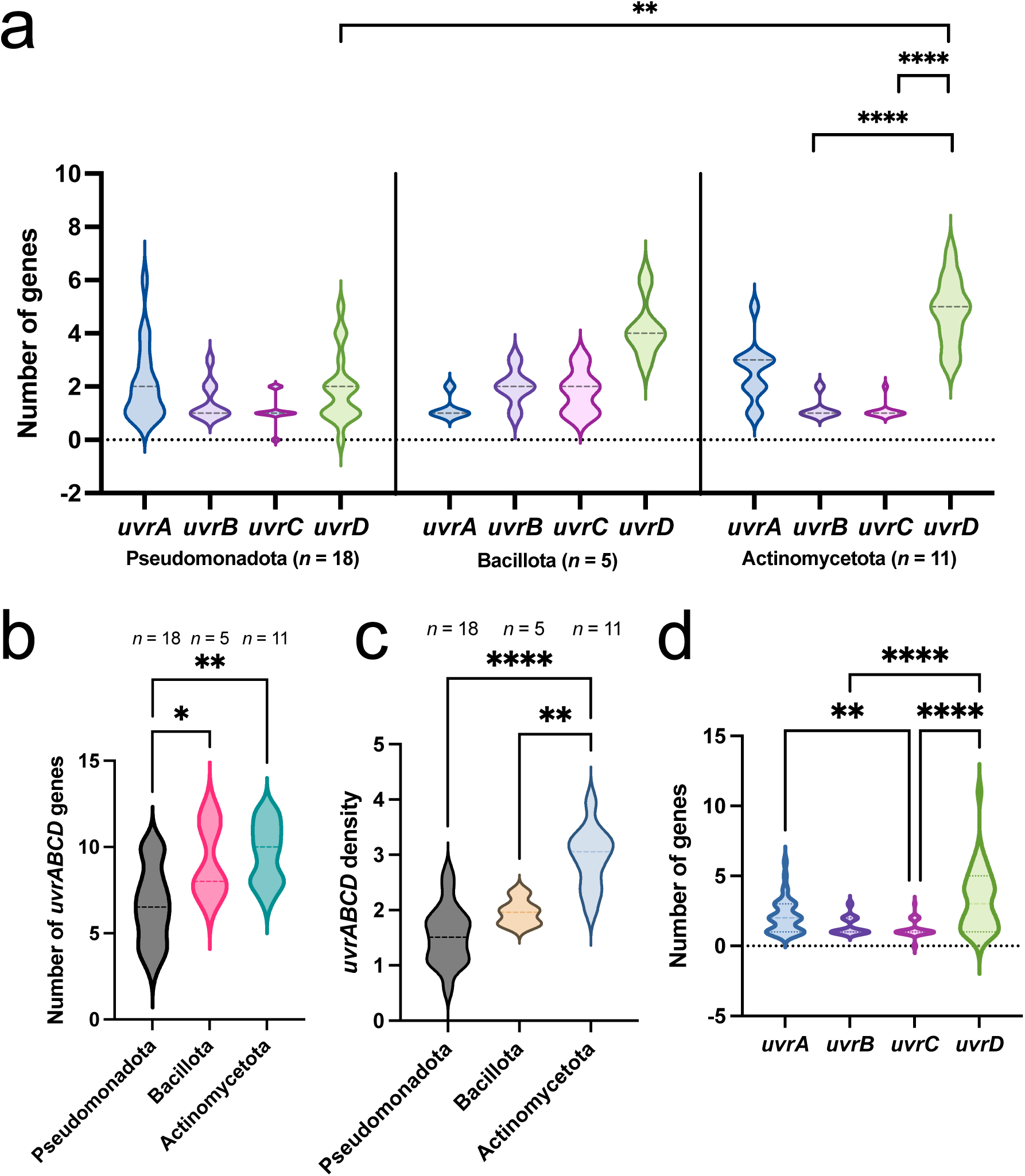
**Distribution and comparison of UVR-encoding genes in the cloud water bacteria**. **a,** The phylum Actinomycetota encoded a significant higher number of *uvrD* as compared to *uvrB* and *uvrC* and compared to *uvrD* of Pseudomonadota. Only significant results within phyla and between the same genes across phyla have been highlighted. **b,** The uvrABCD genes were encoded significantly more in Actinomycetota and Bacillota than in Pseudomonadota. **c,** The *uvrABCD* gene density per bp of the genome is significantly higher in Actinomycetota as compared to Pseudomonadota. Only significant results have been highlighted. **d,** Across all phyla (*n* = 38 genomes), the number of *uvrD* genes was significantly more abundant than *uvrB* and *uvrC* (Kruskal-Wallis test, *p* < 0.0001). *UvrC* was significantly more abundant than *uvrA*. Dashed lines indicate the median. All statistical results can be found in Supplementary Table S8.

Functional annotations of viral-encoded ORFs indicated several genes that could influence viral and host survival in the clouds, such as a *cspA* cold shock protein (KEGG db, <u>K03704</u>) found on two vOTUs, osmotically inducible lipoprotein *OsmB* (<u>K04062</u>) detected on four vOTUs, sensors of blue-light using FAD (Pfam db, <u>PF04940.18</u>) on two vOTUs, ultra-violet resistance protein B (<u>PF12344.14</u>) on two vOTUs, high light inducible protein (VogDB, VOG39308) on one vOTU, impB/mucB/samB family (<u>PF00817.25</u>) involved in UV protection on ten vOTUs. All functional annotations are shown in Supplementary Table 9).

### iRep predicts active replication of cloud water bacteria

To assess the activity of isolates based on their genomic data, iRep values were calculated (Fig. 5d, Supplementary Table 10). iRep estimates microbial replication rates by quantifying genome coverage gradients between the replication origin and terminus. iRep values between 1 and 2 indicating active DNA replication were found in a *Chitinophagaceae* bacterium (MAG01), Candidatus *Novosphingobium monteverdense* (MAG11), Candidatus *Ralstonia monteverdensis* (MAG12), a *Xanthobacteraceae* bacterium (MAG13), *S. melonis* (MAG14), Candidatus *Duganella nubigena* (MAG23), Candidatus *Sphingomonas nubigena* (MAG25), *Acidimicrobiia* bacterium (MAG32), *Aquabacterium* sp. (MAG35), Candidatus *Deinococcus monteverdensis* (MAG38), *Novosphingobium* sp. (MAG41), Candidatus *Variovorax nubigenum* (MAG45), Candidatus *Noviherbaspirillum nubigenum* (MAG46), Candidatus *Methylobacterium nubigenum* (MAG50), *Nocardioides kribbensis* (MAG51), *Sphingomonas paucimobilis* (MAG54), *Pseudarthrobacter* sp. (MAG57), *Agromyces* sp. (MAG59), Candidatus *Pantoea nubigena* (MAG62), as well as isolate genomes *D. nubigenus* MPC36 and *S. panni* MPC37. In addition, iRep values > 2 suggesting that most cells are actively replicating were detected in Candidatus *Spirosoma nubigenum* (MAG55) and the isolate *Curtobacterium nubigenum* MPC39 (Fig. 5d).

### UV-B effects on colony growth and viability of *Curtobacterium nubigenum* MPC39

We explored the UV-B resilience of *C. nubigenum* MPC39, a cloud-borne, rod-shaped bacterium that quickly forms biofilms in liquid culture (Fig. 9g-i). UV-B treatment resulted in no observable change in colony color or morphology of *C. nubigenum* MPC39 over 48 h (Fig. 9a). Colony diameter measurements after four days of incubation showed no significant differences between UV exposure treatments ranging from 0-48 h, although an increasing trend was observed at 12 h exposure, followed by a mild reduction at 24 and 48 h exposure (Fig. 9c). Live/dead staining revealed that all UV treatments significantly increased total cell counts compared to the 0 h baseline (130.4 ± 28.16 cells per microscopic field; ANOVA with Tukey HSD *post hoc*, p < 0.0002), averaging 181.9 ± 76.07, 215.6 ± 64.07, and 514.6 ± 140.25 cells for 12 h, 24 h and 48 h treatments, respectively (Fig. 9b & 9d). Live cell counts followed a similar pattern to colony diameter, peaking at 12 h exposure with a 19.7 ± 54.9 % increase relative to baseline, then declining moderately at 24 h (10.5 ± 12.8 %) and 48 h (2.9 ± 58.7 %), though all timepoints remained elevated to the 0 h baseline (27.9 ± 13.3 %) (Fig. 9d). These results indicate that *C. nubigenum* MPC39 exhibits substantial UV-B tolerance, maintaining viability and colony-forming ability throughout extended UV-B exposure despite fluctuations in cell counts over time.

**Figure 9:**
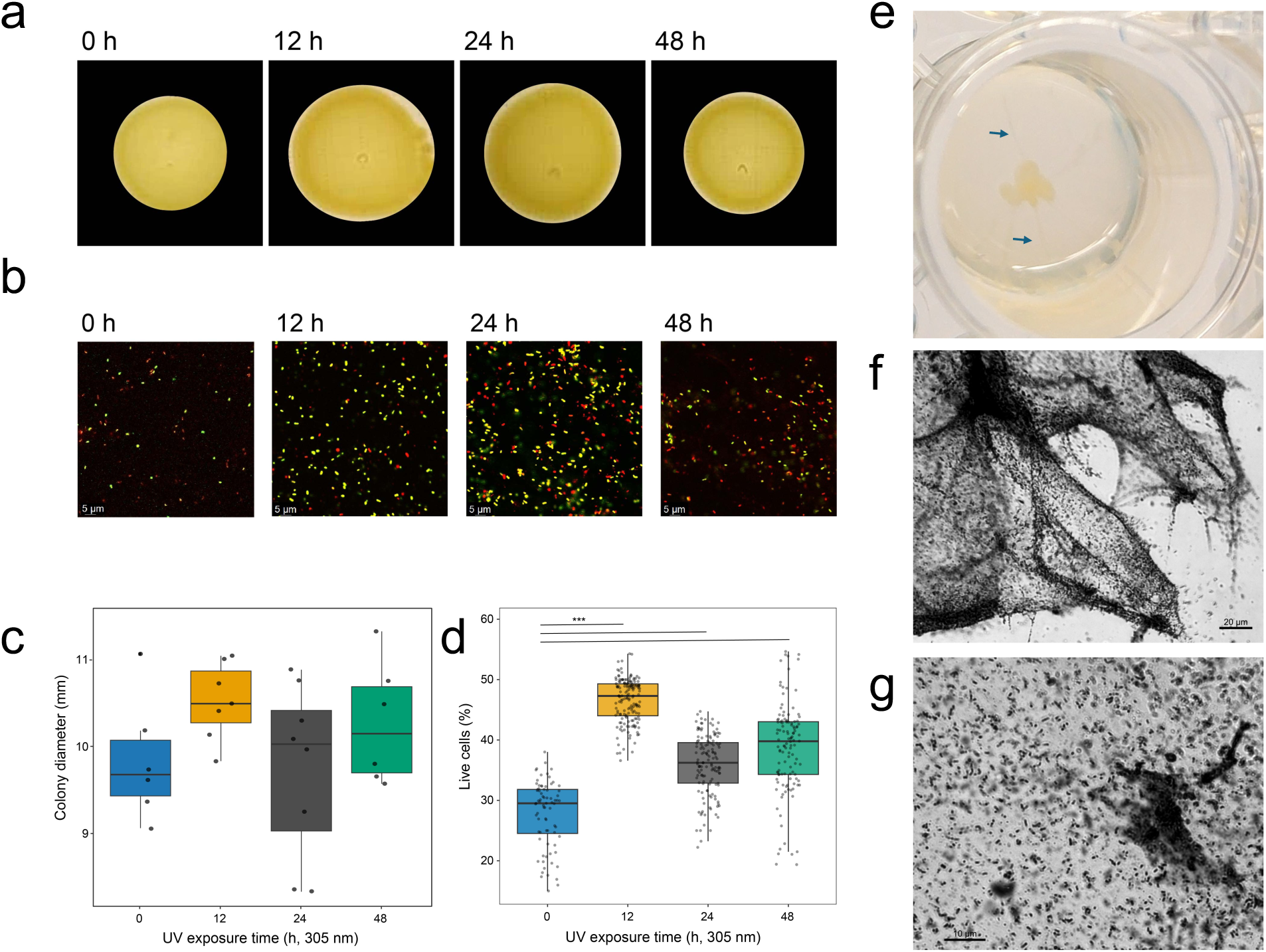
UV-B resistance of *Curtobacterium nubigenum* MPC39 and induction of its prophage. a, Representative colony morphology following UV-B exposure (305 nm) for 0, 12, 24 and 48 h. **b,** Confocal microscopy overlayed images showing live/dead cell distribution after UV exposure (63 × magnification, confocal microscope) for 0, 12, 24 and 48 h. **c,** Colony diameter measurements across treatments (*n* = 5 - 6 per treatment). **d,** Quantification of viable and non-viable cells from fluorescence imaging (*n* = 4 - 5 per treatment). **e,** *C. nubigenum* MPC39 forms a biofilm in the well of a microtiter plate with appendages attaching to the well wall (blue arrows). **f,** Biofilm of *C. nubigenum* MPC39. **g,** The strain is rod-shaped in light microscopy.

### Prophage induction for cloud water strain *C. nubigenum* MPC39 and taxonomic classification

Since no lytic phages were identified, we tested whether one or both prophages detected in the genome of the cloud water strain *C. nubigenum* MPC39 could be induced into the lytic cycle. Both were predicted as intact by PHASTEST (Fig. 10a). Prophage 1 (40 kb size, 68.4 % G+C content) showed 10 % intergenomic similarity and 59.9 % mean protein identity (over 20 % genome length, tBLASTx) to Microbacterium phage Footloose (Actinomycetota host group). Prophage 2 (48.1 kb size, 49.9 % G+C content) and was identical (in sequence and length) to Escherichia phage Lambda (Pseudomonadota host group) (Fig. 10e). Mitomycin C treatment at 0.5 and 1 µg mL^-1^ triggered prophage 1 (confirmed by sequencing of supernatant DNA; Fig. 10e), as shown by a decline in *C. nubigenum* MPC39 growth after four hours (Fig. 10b). Transmission electron microscopy (TEM) revealed siphovirus particles in the supernatant, confirming induction (Fig. 10c). We propose the name Curtobacterium phage vB_CnuS_Cirrus1 for this novel phage. It contains 63 ORFs and a serine tRNA (Fig. 10d; Supplementary Table S11). Sharing 11 core proteins with Microbacterium phage Footloose (Supplementary Table S12), it likely belongs to the same currently unnamed viral family, for which we propose the name Nebulaviridae. We further propose the genus name “Nebulavirus” and the species name “Nebulavirus nubigenus”. Assuming that all phages observed in TEM belong to one type, capsid and tail dimensions were measured as 51.2 ± 3.8 nm and 139.1 ± 10.9 nm, respectively (Supplementary Table S13). No prophage induction occurred after UV-B exposure, as hourly OD_600_ measurements showed recovery in all cultures (Supplementary Fig. S11). These results indicate that at least one prophage in *C. nubigenum* MPC39 is inducible by mitomycin C but not by UV-B, highlighting selective triggers for prophage activation in this cloud water bacterium.

**Figure 10:**
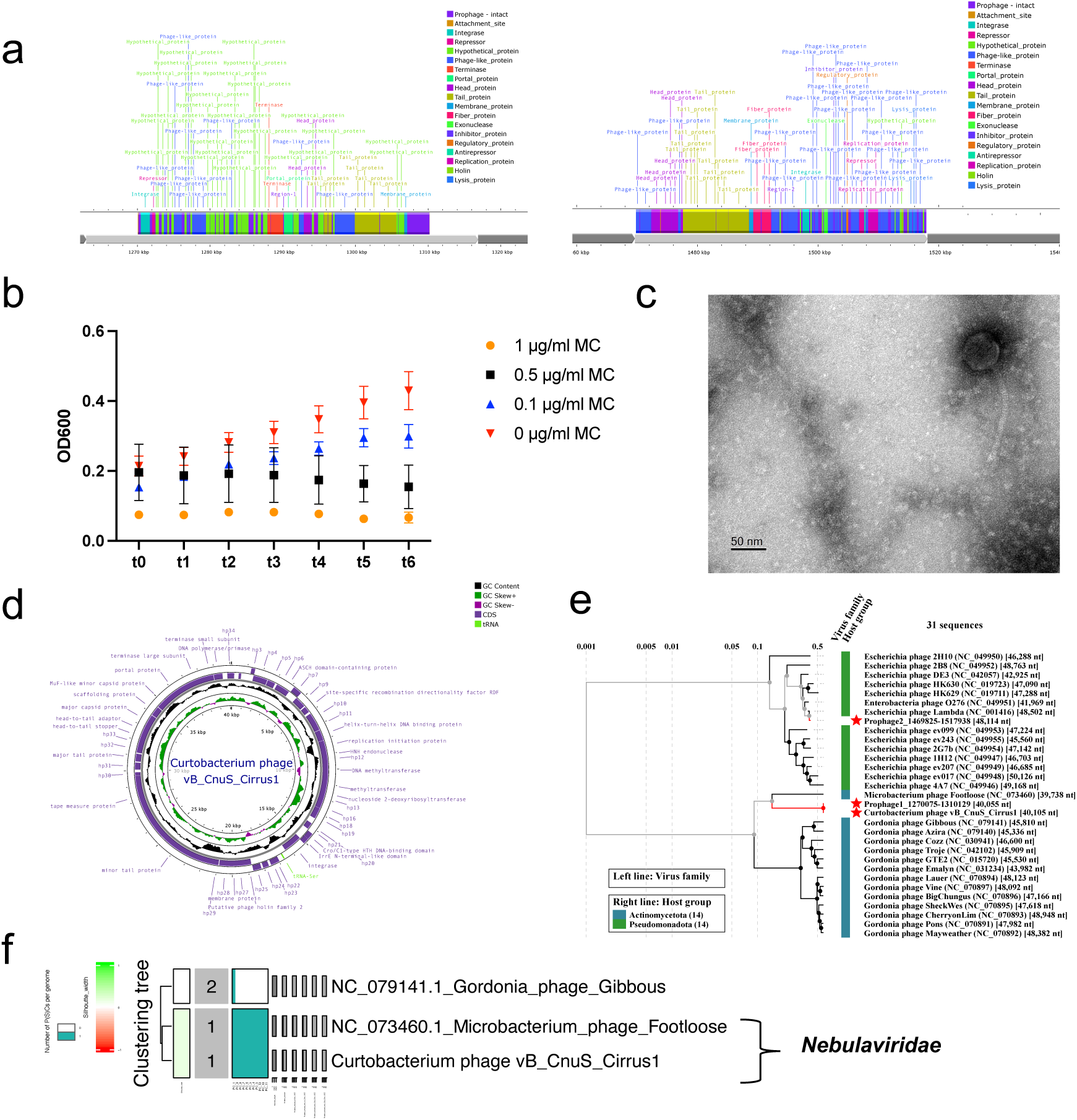
Induction of a prophage from *Curtobacterium nubigenum* MPC39. a, Genomic sequences of two prophages within the genome of *C. nubigenum* MPC39 predicted by PHASTEST. **b,** Growth curve of the strain under different mitomycin C concentrations. **c,** Transmission electron microscopy image of a *Curtobacterium* phage from the supernatant after induction. **d,** Circularized genome of Curtobacterium phage vB_CnuS_Cirrus1 with functional annotations. **e,** Identity of the Curtobacterium phage vB_CnuS_Cirrus1 was revealed by sequencing DNA from the assays’ supernatant and protein-based alignment with the two prophages of the host (Prophage1 and Prophage2). The tree was generated using ViPTree v.4 [75]. **f,** Curtobacterium phage vB_CnuS_Cirrus1 forms the family *Nebulaviridae* together with Microbacterium phage Footloose.

## Discussion

Our study provides a comprehensive examination of viral diversity in cloud water, revealing the presence of active prophages, multiple viral variants across proximate cloud events, CRISPR spacer-protospacer interactions, and a broad range of predicted bacterial hosts, many of which represent previously unknown species. Our findings underscore that cloud water is not merely a passive collection of microorganisms transported from terrestrial and aquatic sources, but a dynamic environment in which viral-host interactions are actively shaping microbial community structure. This is an astonishing finding, given that only one out of > 1000 cloud water droplets are estimated to contain a bacterial cell [8]. The detection of 49 proviruses in the viral metagenome plus active prophages in a cloud water-derived *C. nubigenum* MPC39 suggests that lysogenic phages may contribute significantly to microbial turnover in this ephemeral habitat, potentially influencing bacterial population dynamics through lysis-mediated release of nutrients and horizontal gene transfer. Prophages have high prevalence in atmospheric bacteria [83], and environmental conditions such as high UV radiation can trigger their induction [84]. Lysogens may also exhibit greater resistance to UV light than non-lysogenic counterparts [84]. As no phage induction was observed during UV-B exposure, these findings support the idea that UV resistance in this strain may be associated with lysogeny, which would require further investigation.

The observation of multiple viral variants (single nucleotide variants) in cloud water samples collected within a few days indicates rapid turnover and diversification within cloud-associated viral populations, suggesting that these communities are highly dynamic and may respond quickly to environmental changes or stochastic dispersal events. The presence of CRISPR arrays with spacer to viral protospacer matches in bacterial strains that commonly occur in cloud water, like the genera *Sphingomonas* and *Deinococcus* [85, 86], demonstrates that bacterial populations in clouds are actively responding to viral predation and maintain adaptive immunity, which indicates that virus-host coevolution in these systems extends beyond transient cloud events. While some interactions could theoretically have occurred after sampling within the collection bottles, the CRISPR evidence points to longer-term evolutionary dynamics, consistent with repeated or historical encounters between hosts and phages rather than being solely a product of post-sampling activity.

We found predicted bacterial hosts for cloud-borne viral populations spanning multiple taxa which further emphasizes the ecological breadth of cloud-associated viruses. The presence of viruses infecting both common atmospheric bacteria and more specialized taxa highlights the potential for viruses to mediate microbial diversity and functional potential. Given that cloud water can influence cloud chemistry and potentially impact precipitation, viral lysis and microbial metabolism may have indirect biogeochemical consequences, for example by releasing organic carbon or other compounds that alter cloud droplet chemistry or aerosol interactions. We found that bacteria present in cloud water like *Deinococcus* spp. and *Cand.* Ralstonia monteverdensis carried genes encoding carbon monoxide dehydrogenase (CODH) units, suggesting that these microorganisms may play a role in atmospheric CO cycling. CODH is an enzyme complex that catalyzes the oxidation of CO to CO_2_, providing a source of energy and electrons for microbial metabolism. This parallels findings in aerobic soil actinobacterium *Mycobacterium smegmatis*, where atmospheric trace gases have been proposed to sustain ATP production under nutrient-limited conditions, and to enhance upregulated expression of form I molybdenum-copper carbon monoxide dehydrogenase [87, 88]. The presence of CODH genes in cloud microbiota indicates that CO may serve as a “lifeline” energy substrate, enabling bacteria to maintain basal metabolism and proton-motive force even in the oligotrophic and oxidative environment of cloud droplets. This is consistent with studies showing that microbes often rely on atmospheric trace gases, e.g., H_2_, CO, CH_4_, as alternative energy sources in nutrient-poor environments [89]. Osmoregulatory genes were present in multiple genomes, likely reflecting that the cloud environment is highly variable in water and solute content. These genes, probably often inherited from their original habitats such as leaves and soil surfaces, allow the bacteria to withstand osmotic stress in cloud droplets and help them survive long enough for transport and deposition. We also detected *Curtobacterium* and *Deinococcus* in our samples. In other (atmospheric) environments, these genera are well known for their resilience, with *Deinococcus radiodurans* exhibiting resistance to ionizing and UV radiation, desiccation, and oxidative stress [90–92]. *Curtobacterium aetherium*, previously isolated from the stratosphere [93, 94], is also highly UV radiation resistant. In the present study, the moderate reduction in colony growth, with minimal loss in live cell proportions, and no morphological alterations, indicates a high level of resilience to UV-B exposure of *C. nubigenum* MPC39. Together with the high iRep value indicating active replication, this suggests that this isolate is well adapted to the UV-stressed conditions characteristic of cloud environments, supporting its ecological persistence in atmospheric habitats. The presence of multiple CSP and UV irradiation genes in multiple genomes and some vOTUs indicates significant environmental adaptations, such as to fluctuating temperatures and constant UV stress of cloud environments. Additionally, the abundance of uvrABCD, suggests the demand for enhanced SOS and DNA repair mechanisms in response to UV stress, particularly uvrD, which performs an essential role in maintaining genomic stability and DNA lesion repair [95].

A limitation of our study lies in the sampling protocol. Between cloud water collection events, the sampler was cleaned with ultrapure water, without the use of sterilizing agents such as bleach or ethanol, or an autoclaving protocol, which was not logistically feasible. This introduces the possibility of carryover or cross-contamination, which could artificially inflate observed viral diversity or obscure temporal patterns. While the repeated detection of CRISPR spacer matches, consistent host predictions, and ecological plausibility of the viral assemblages suggest that most detected virus-host interactions are biologically meaningful, future studies should implement more stringent sterilization methods of the cloud water collectors to minimize potential artifacts. Furthermore, the samples were stored frozen at -20 °C for seven years prior to processing, which may have biased the isolated fraction toward bacteria capable of surviving prolonged freeze-thaw cycles and those carrying cold shock genes. On the other hand, this may reflect natural selection pressures in clouds, where microorganisms frequently experience freezing and desiccation [96, 97], suggesting that the freeze-tolerant taxa we detected are ecologically relevant.

Another consideration is the short residence time and highly variable nature of cloud water. Microbial communities in clouds are influenced by numerous factors, including atmospheric mixing, precipitation events, and deposition from terrestrial and aquatic sources. Emissions of viruses from soils and oceans are expected [98, 99], or, alternatively, viruses travel within surface-emitted bacteria. Thus, while the detection of active prophages and CRISPR interactions points to ongoing ecological processes, these processes occur on very short timescales, and the relative importance of local versus long-range sources remains uncertain. Future high-frequency temporal studies, ideally combined with metatranscriptomic or single-virus approaches, could help disentangle contributions of in-cloud activity and long-range or short-range dispersal. Future work should aim to resolve the temporal scale of these interactions and explore the mechanistic consequences of viral activity in clouds, including potential impacts on microbial metabolism and atmospheric biogeochemical cycling.

Our work has several important implications. Because clouds are extreme, dynamic environments analogous to extraterrestrial atmospheres, the stress-protective features that allow Earth’s cloud microbes to survive and disperse provide direct insight into how life could adapt, persist, and spread on other planets [100]. By revealing predicted virus-host interactions, viral variants, stress-resistance genes, a UV-tolerant isolate with active prophages, including several previously unrecognized bacterial strains, our study expands the known functional and taxonomic diversity of cloud microbiomes. These findings highlight clouds as dynamic reservoirs of microbial and viral innovation, with implications for atmospheric dispersal, ecosystem connectivity, and the potential for life to withstand extreme conditions on Earth and beyond.

## Conclusions

Taken together, our results provide strong evidence that cloud water harbors active viral populations with ongoing virus-host interactions, reinforcing the concept of clouds as ephemeral yet ecologically significant microbial and viral habitats. These findings expand current understanding of viral ecology in atmospheric systems and suggest that viral dynamics should be considered alongside bacterial diversity in studies of atmospheric microbial biogeography and cloud biogeochemistry. Importantly, while some interactions may have occurred after sampling, the CRISPR signatures indicate longer-term virus-host coevolution, reflecting evolutionary processes that likely extend across multiple cloud cycles.

## Supporting information

Register List Seqcode

## List of abbreviations

ANOVA: Analysis Of Variance
BLAST: Basic Local Alignment Search Tool
CO: carbon monoxide
CSP: cold shock protein
CRISPR: Clustered Regularly Interspaced Short Palindromic Repeats
DNA: deoxyribonucleic acid
G+C: guanine-cytosine
iRep: index of replication
KEGG: Kyoto Encyclopedia of Genes and Genomes
KO: KEGG Orthology
MAG: metagenome-assembled genome
MC: mitomycin C
TEM: transmission electron microscopy
OD: optical density
ORF: open reading frame
UV-B: ultraviolet B
VC: viral cluster
vOTU: viral operational taxonomic unit

## Declarations

## Ethics approval and consent to participate

Not applicable

## Consent for publication

Consent for publication was provided for Figure 1.

## Availability of data and material

The sequencing datasets supporting the conclusions of this study are available in the NCBI Sequence Read Archive (SRA) under the corresponding BioProject ID PRJNA1273110 and BioSample accessions SAMN48932728 - SAMN48932734. The 24 bacterial isolate genomes have been deposited in GenBank under BioSample accession numbers SAMN48997597 - SAMN48997620. The 62 metagenome-assembled genomes (MAGs) are also available under BioSample accessions SAMN48999342 – SAMN48999403. The viral operational taxonomic units (vOTUs) are stored at GenBank under accessions PV837008 - PV837465. For detailed information, including run accession numbers, please refer to Supplementary Table S15. TEM images are accessible via Figshare at doi: 10.6084/m9.figshare.30354244. FLEXPART code is available from flexpart.eu.

## Authors’ contributions

Conceptualization: J.R., N.LY., and M.v.P. Methodology: J.R., N.LY., E.L., J.M. Investigation: J.R., N.LY., E.L., G.W., R.D., K.B., R.B., S.E., N.E., C.G.Z. Visualization: J.R., N.LY., E.L., G.W., R.D., K.B., S.E., N.E., C.G.Z. Funding acquisition: J.R., M.v.P. Project administration: J.R., M.v.P, Writing-original draft: J.R., G.W. Writing-review and editing: all co-authors.

## Acknowledgements

We like to thank Pierre Amato for discussions about viruses in clouds. We acknowledge technical assistance by Bingli Clark Chai during cloud water processing. We further acknowledge the use of the HPC cluster DRACO, with tools and database partly provided by the VEO group of Bas Dutilh at the FSU Jena. We also thank the PDC Center for High Performance Computing, KTH Royal Institute of Technology, Sweden, for providing access to the computing resources used in this research (NAISS 2025/22-272, NAISS 2025/23-14). We thank the Core Facility for next-generation sequencing of the Leibniz Institute on Aging-Fritz Lipmann Institute in Jena for Illumina sequencing. Here, we acknowledge Muriel Ritsch, Ivonne Görlich, and Marco Groth for their support.

