## Supplementary material for "Virus-host interactions and viral population dynamics across atmospheric cloud events"

**New bacterial species from cloud water**

*Curtobacterium* sp. MPC1 (99.99 % completeness in CheckM2) had the highest average nucleotide identity based on BLAST (ANIb) to *Curtobacterium luteum* NS184 (91.66 % ANIb, 76.7 % aligned fraction), and hence we propose the new species name *Curtobacterium capverdeinubigenum* MPC1 (capverdi for its origin from the Cape Verde islands, Latin *nubes*, cloud; *-genus*, born of).

*Curtobacterium* sp. MPC39 (100 % completeness) had 98.22 % ANIb, 78.49 % aligned fraction to *Curtobacterium* sp. B18 (Genbank, GCF_000333375.1). Hence it is the same genomic species, but since the clade lacks a formal name, we propose the name *Curtobacterium nubigenum* MPC39 for this strain. There was another isolate of *Curtobacterium citreum* MPC3 (100 % completeness) with 98.69 % ANIb and 84.72 % aligned fraction to the reference genome *Curtobacterium citreum* NS330, hence we assigned this name. In addition, two new *Deinococcus* species were found. Candidatus *Deinococcus monteverdensis* MAG38 (94.69 % completeness, the name pointing towards its origin from Mont Verde) had 80.93 % ANIb and 65.37 % aligned fraction to the next representative *Deinococcus* sp. KSM4-11. Another cloud water isolate named *Deinococcus nubigenus* MPC36 (100% completeness) had 91.11 % ANIb and 66.67 % aligned fraction to the known strain *Deinococcus gobiensis* I-0. The MAG and the isolate genome of the two cloud-water derived *Deinococcus* had only 76.4 % ANIb of 34 % aligned fraction to each other.

We propose the name *Agrococcus nubigenus* MPC7b for another isolate (99.98 % completeness), which had 90.51 % ANIb and 71.51 % aligned fraction to the type strain *Agrococcus jejuensis* DSM 22002, which was previously isolated from dried seaweed. Candidatus *Ralstonia monteverdensis* is the proposed name for MAG05 (99.88 % completeness), which had 99.63 % ANIb and 94.97 % aligned fraction to *Ralstonia* sp. MD27, and 93.32% ANIb and 81.55 % aligned fraction to *Ralstonia insidiosa* ATCC 49129. Candidatus *Novosphingobium monteverdense* MAG11 (98.75 % completeness) had 85.77 % ANIb and 72.27 % aligned fraction to *Novosphingobium sediminicola* DSM 27057 and thus represents a new species. The MAG Candidatus *Variovorax nubigenus* MAG45 (89.55 % completeness) had 77.59 % ANIb and 51.5 % aligned fraction to *Variovorax beijingensis* 502. A MAG of proposed Candidatus *Duganella nubigena* MAG23 (99.99 % completeness) shared 97.71 % ANIb and 89.72 % aligned fraction with *Duganella* sp. Leaf126 as well as 86.39 % ANIb and 71.96 % aligned fraction with *Duganella zoogloeoides* ATCC 25935.

Candidatus *Noviherbaspirillum nubigenum* MAG46 (92.92 % completeness) was assigned a new species name as it shared only 88.59 % ANIb and 71.96 % aligned fraction with *Noviherbaspirillum soli* SUEMI10. The newly proposed Candidatus *Methylobacterium nubigenum* MAG50 (93.88 % completeness) had 89.23% ANIb and 77.72% aligned fraction to *Methylobacterium phyllostachyos* BL47. Candidatus *Pantoea nubigena* MAG62 (95.52 % completeness) had 73.4 % ANIb and 58.39 % aligned fraction to *Pantoea vagans* ND02 and thus also at the outer edge of the *Pantoea* genus but certainly a new species as proposed. Candidatus *Spirosoma nubigenum* MAG55 (90.19 % completeness) had 83.27 % ANIb and 81.2 % aligned fraction to *Spirosoma rhododendri* CJU-R4 and thus represents a new species.
Candidatus *Sphingomonas nubigena* MAG25 (99.61 % completeness) had 87.4 % ANIb and 76.52 % aligned fraction to *Sphingomonas palmae* JS21-1 and thus received this new species name. *Sphingomonas* sp. MAG14 had 98.7 % ANIb and 86.89 % aligned fraction to *Sphingomonas melonis* and was assigned this name. Here it turned out that *Sphingomonas aquatilis and Sphingomonas melonis* have previously received different names for the same species, since *Sphingomonas melonis* 30a has 95.24 % ANIb to *Sphingomonas aquatilis* DSM 15581. The genome of our isolate *Sphingomonas* sp. MPC37 (99.91 % completeness) had 96.37 % ANIb and 83.92% aligned fraction to *Sphingomonas panni* and was thus named as such. Supplementary Table S7 provides further information about the comparisons described here.

**Plaque Assays**

Plaque assays were performed to screen for bacteriophages in six cloud water samples (WW3, WW8, WW13, WW16, WW25, and WW29) using the following bacterial host strains: *Curtobacterium capverdeinubigenum* MPC1, *Curtobacterium* sp. MPC2, Alkalihalobacillus_A gibsonii_A MPC5 and MPC9, and *Micrococcus luteus* MPC11. MPC1 and MPC2 were grown on R2A agar, while the remaining strains were grown on LB. Overnight bacterial cultures were replenished with 1 mL of fresh medium and incubated for an additional hour at 135 rpm. For each assay, 200 μL of cloud water was added to base agar plates, followed by overlaying with 3.5 mL of MSM top agar (450 mM NaCl (Sigma), 50 mM $MgSO_4$ x 7 $H_2O$ (Sigma), 50 mM Trizma base (Sigma), 5 g $L^{-1}$ low-melting agarose) containing 300 μL of the respective bacterial culture. Plates were swirled to ensure even distribution, allowed to solidify, and incubated for 24-48 h to monitor plaque formation. Clear zones observed were picked using sterile pipette tips, resuspended in MSM buffer (top agar without low-melting agarose), centrifuged, and stored at 4 °C. For further purifications, each resuspension in MSM buffer obtained was streaked onto fresh top agar plates with the appropriate bacterial host and incubated for 24-48 h. All procedures were conducted under sterile conditions. Plaque assays initially revealed some clear zones suggestive of phage activity in multiple cloud water samples across different bacterial hosts. However, these clear zones did not reappear upon purification, suggesting that the initial observations were likely air bubbles rather than true phage plaques.

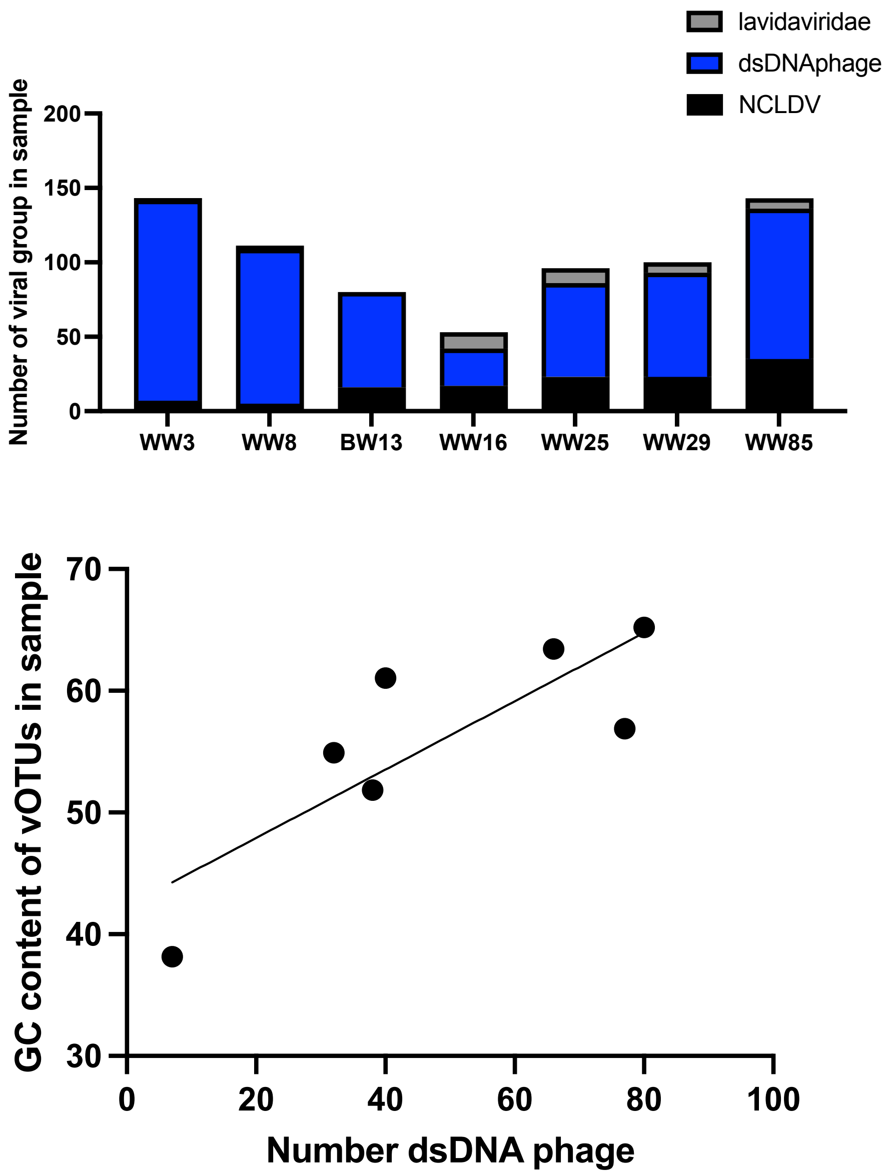


**Figure S1:** Viral groups identified by Virsorter2 in different samples with their presence determined by read mapping (upper panel). A positive correlation (Pearson's $r$= 0.82, $p$ = 0.023, $n$ = 7) could be observed between the GC base content of viral operational taxonomic units (vOTUs) present in a sample and the number of dsDNA phages (for both presence was determined based on read mapping) (lower panel).

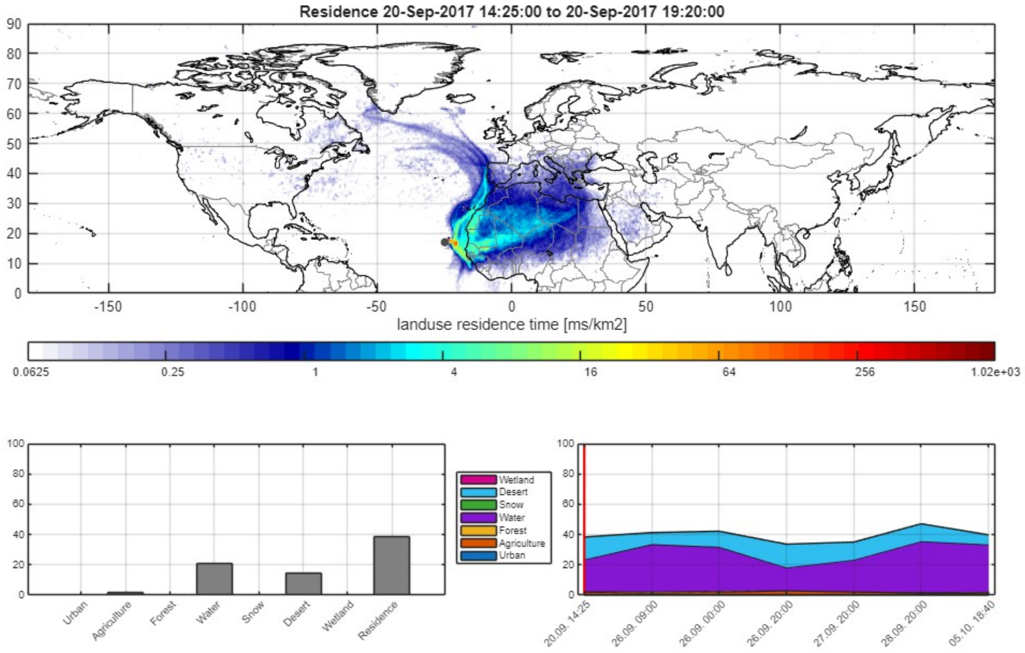


**Figure S2:** Emission sensitivity for the sample starting on the 20.9.2017 14:25, the lower left panel shows the residence time over different land use categories, and the lower right panel shows the land use categories desert and ocean for all samples.

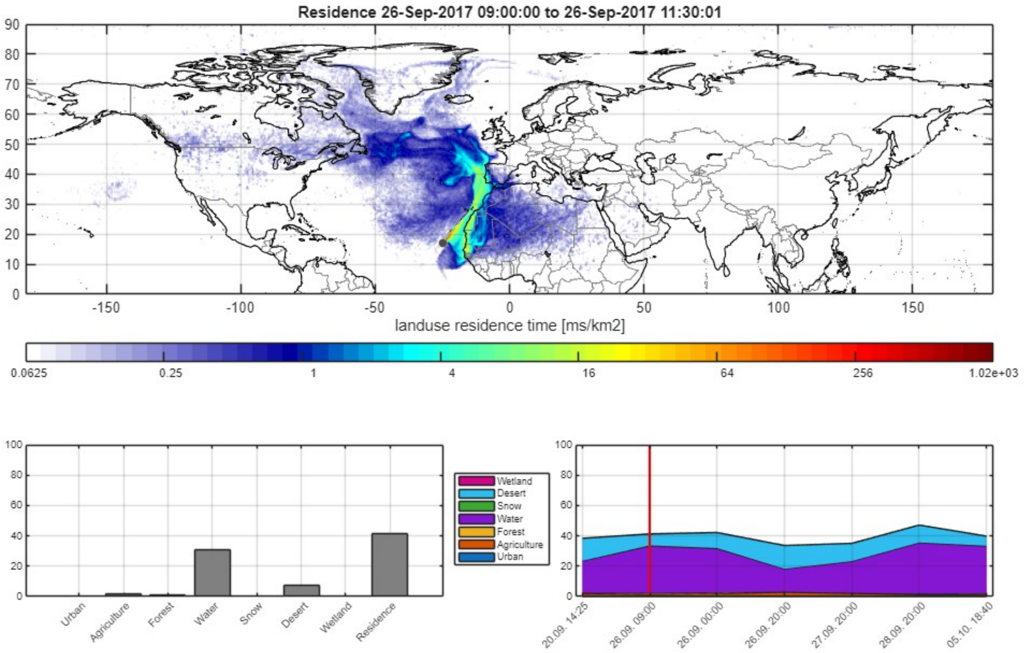


**Figure S3:** Emission sensitivity for the sample starting on the 26.9.2017 09:00, the lower left panel shows the residence time over different land use categories, and the lower right panel shows the land use categories desert and ocean for all samples.

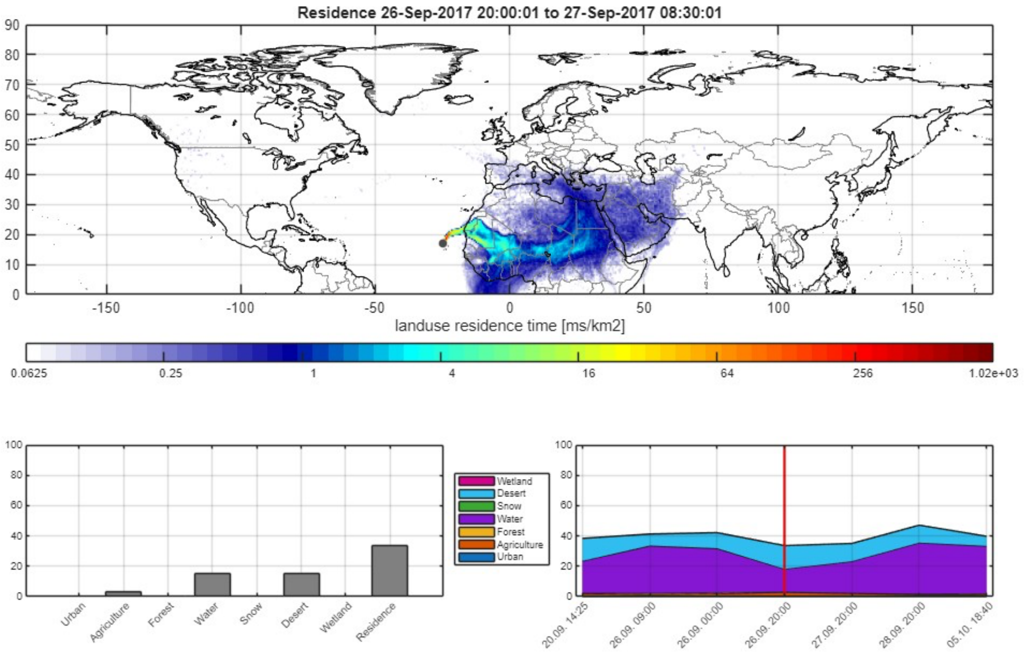


**Figure S4:** Emission sensitivity for the sample starting on the 26.9.2017 20:00, the lower left panel shows the residence time over different land use categories, and the lower right panel shows the land use categories desert and ocean for all samples.

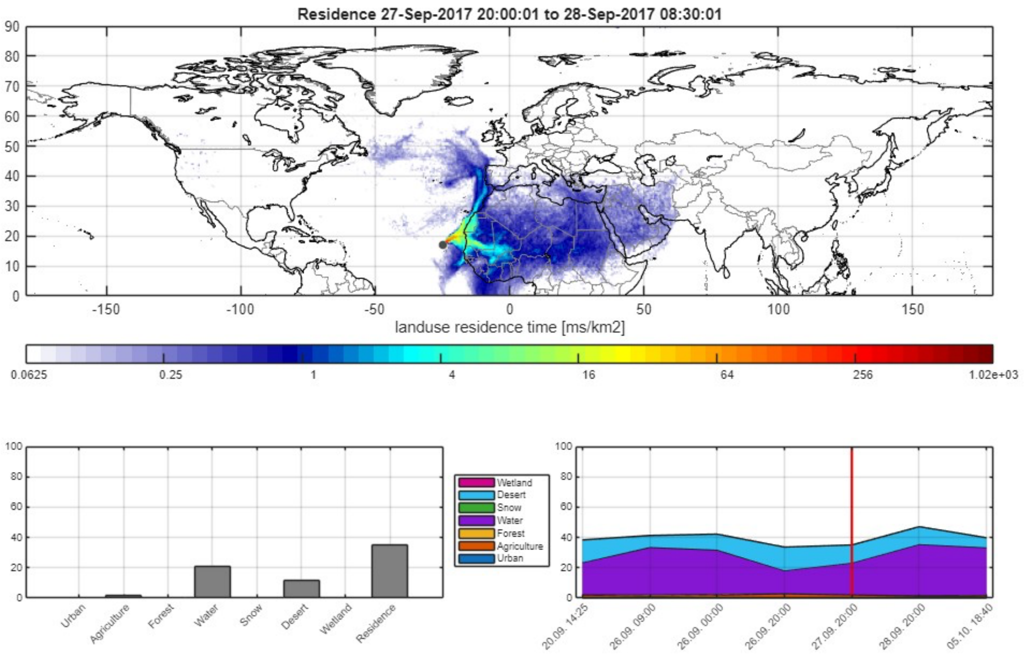


**Figure S5:** Emission sensitivity for the sample starting on the 27.9.2017 20:00, the lower left panel shows the residence time over different land use categories, and the lower right panel shows the land use categories desert and ocean for all samples.

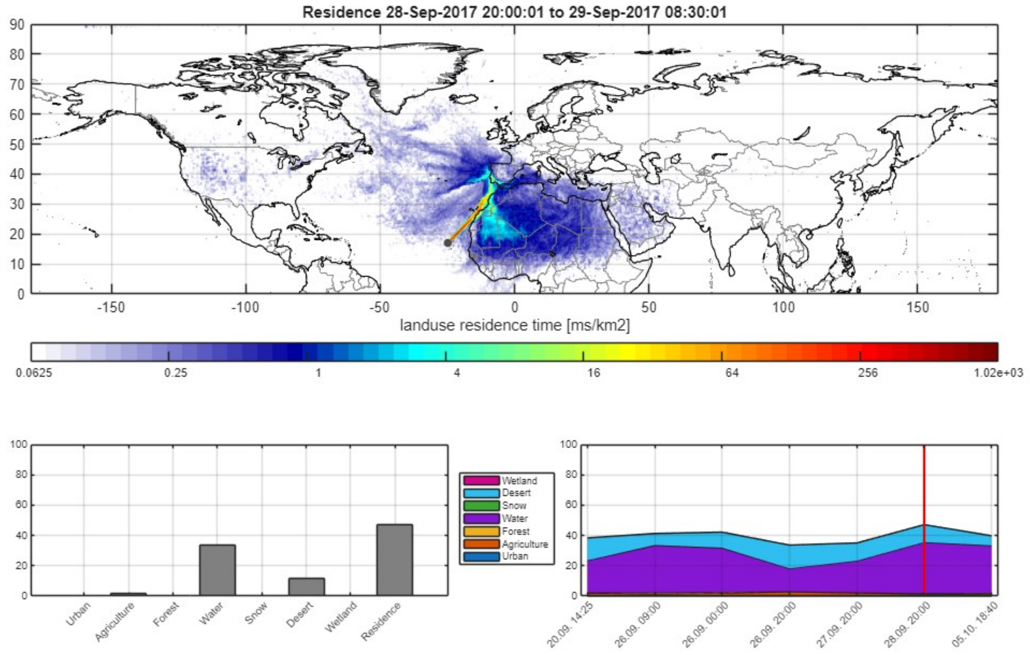


**Figure S6:** Emission sensitivity for the sample starting on the 28.9.2017 20:00, the lower left panel shows the residence time over different land use categories, and the lower right panel shows the land use categories desert and ocean for all samples.

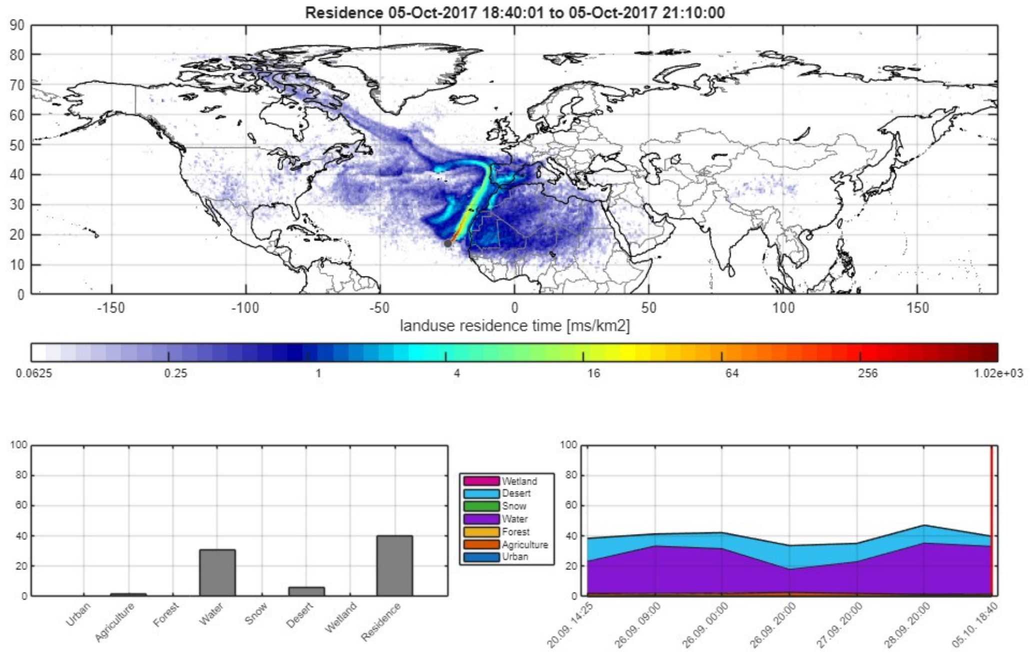


**Figure S7:** Emission sensitivity for the sample starting on the 05.10.2017 18:40, the lower left panel shows the residence time over different land use categories, and the lower right panel shows the land use categories desert and ocean for all samples.

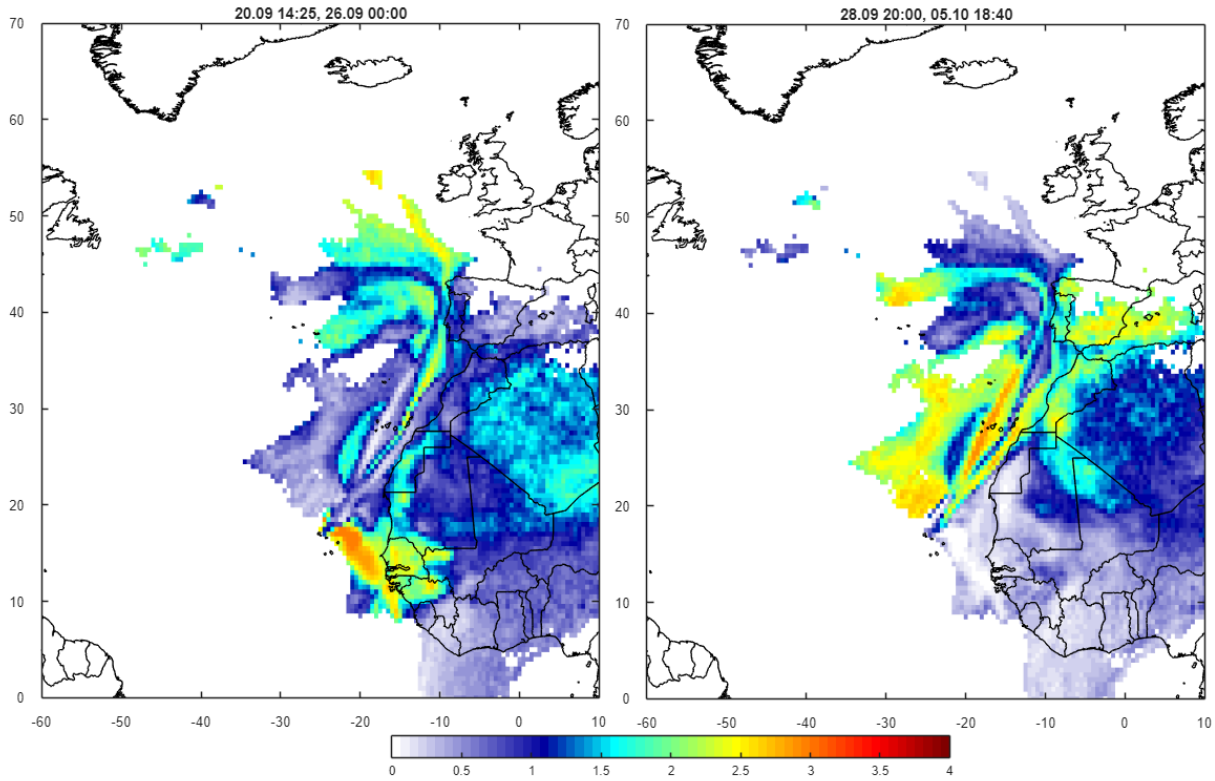


**Figure S8:** Anomaly plot for the 2 episodes. The left (right) panel shows the average emission sensitivities for episode 1 (2) divided by the average emission sensitivity. High values show regions where air masses come from during the episodes, but not during average situations. For episode 1 West Africa is a source region, while for episode 2 Portugal and Spain are source regions.

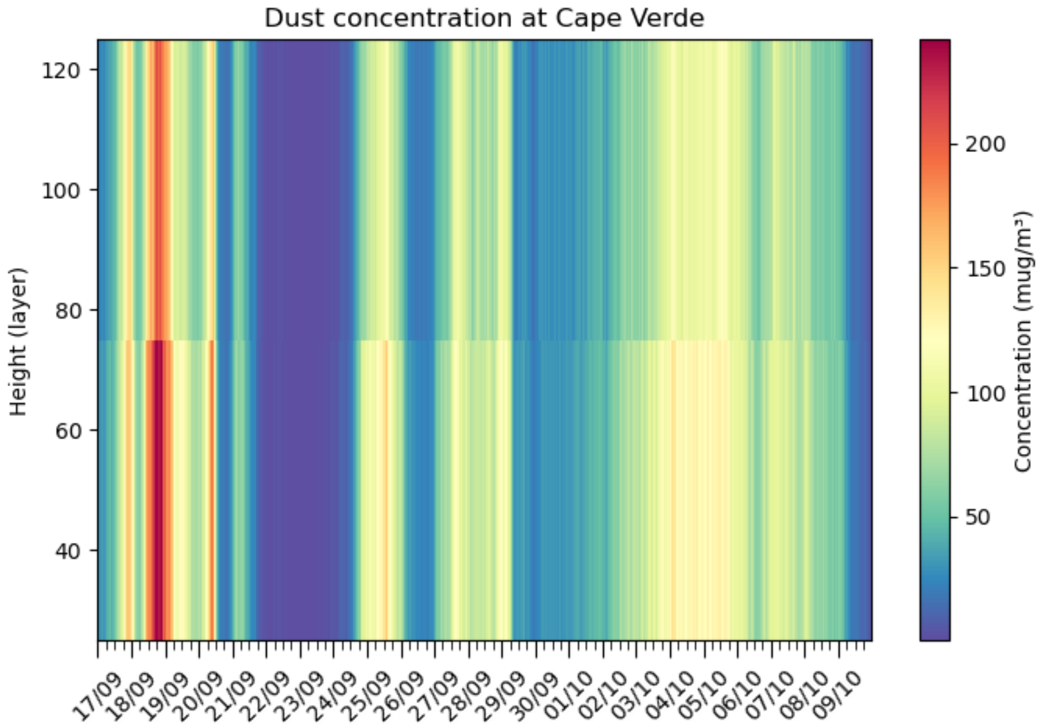


**Figure S9:** FLEXPART simulated dust concentration at Cabo Verde at a height of 50 and 100m.

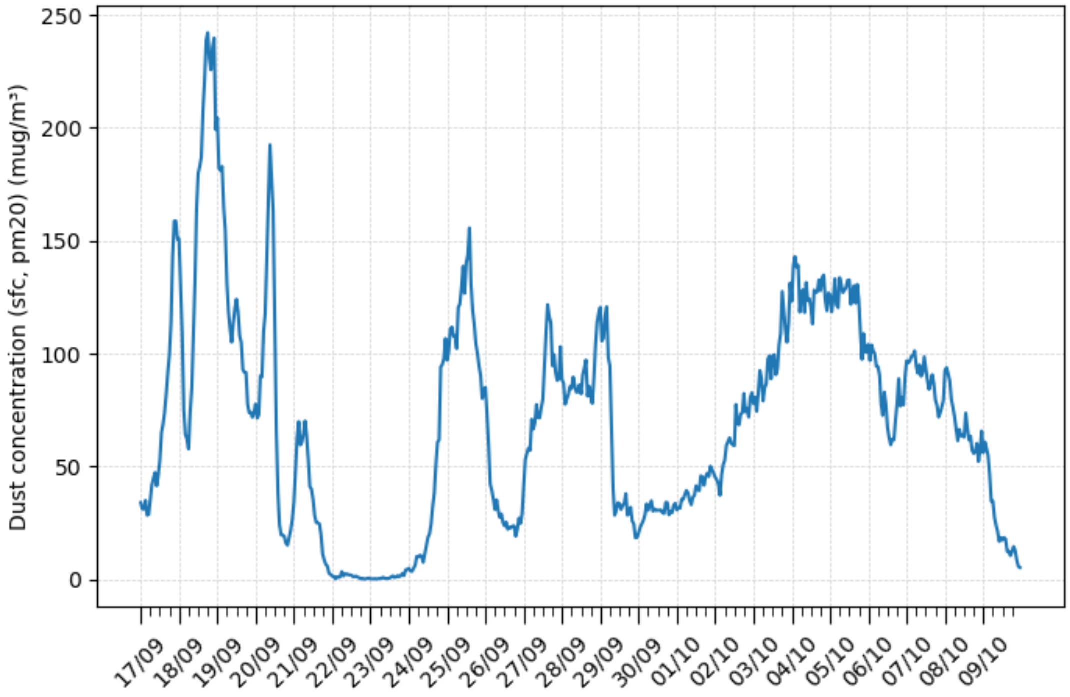

**Figure S10:** FLEXPART simulated surface dust concentration at Cabo Verde, 3 hourly data for the period of 17 Sept. to 9 Oct. 2017.

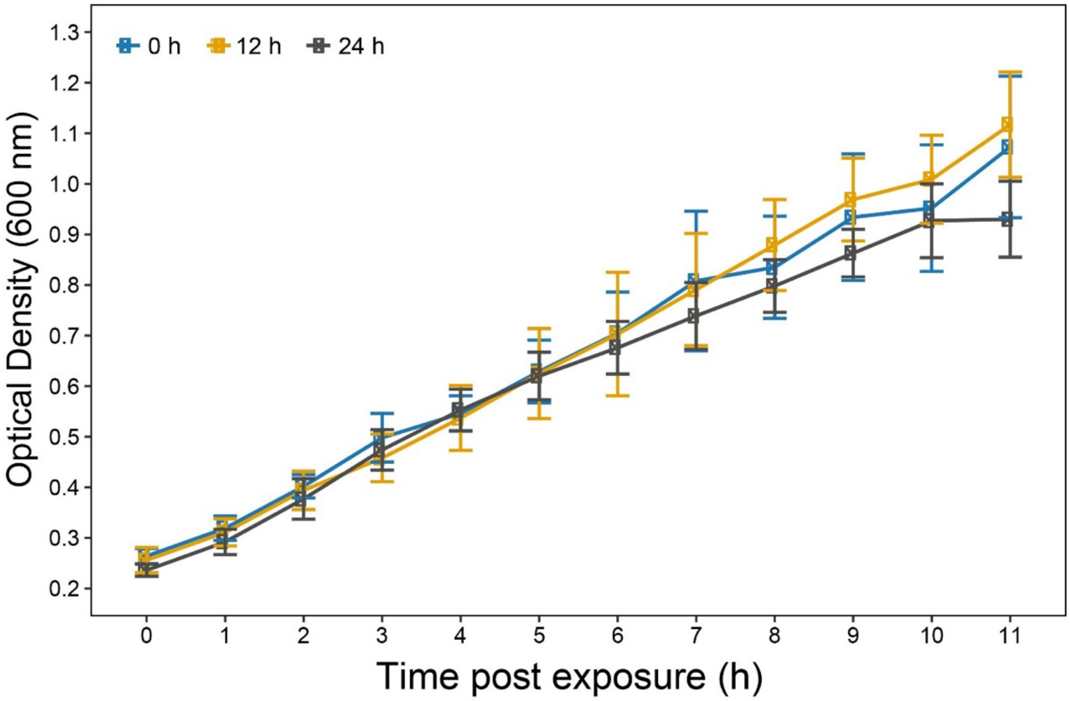


**Figure S11: Recovery growth of *Curtobacterium nubigenum* MPC39 following UV-B exposure.** Growth curves of cultures after exposure to UV-B radiation (305 nm) for 0, 12, or 24 h (n = 4 - 5 per treatment).
