## Supplementary material for "Virus-host interactions and viral population dynamics across atmospheric cloud events": Register List Seqcode

### Register list for 14 new names including Ralstonia monteverdense sp. nov.

**Submitted by Rahlff, Janina**

#### Species *Agrococcus nubigenus*

**Etymology**
[nu.bi.ge'nus] **L. fem. n.** *nubes*, a cloud, mist, vapor; **N.L. masc. adj. suff.** *-genus*, born from; **N.L. masc. adj.** *nubigenus*, born from the clouds

**Nomenclatural type**
INSDC Nucleotide: JBPGPM000000000 Ts

**Description**
*Agrococcus nubigenus* MPC7b grows as red colony, 0.5 mm diameter, round; was isolated from cloud water sample WW85

**Classification**
*Bacteria* » *Actinomycetota* » *Actinomycetes* » *Micrococcales* » *Microbacteriaceae* » *Agrococcus* » *Agrococcus nubigenus*

**Registry URL**
https://seqco.de/i:55992

#### Species *Curtobacterium capeverdeinubigenum*

**Etymology**
[cape.ver.de.i.nu.bi.ge'num] **N.L. fem. n.** *Capeverdea*, referring to Cape Verde; **L. fem. n.** *nubes*, a cloud, mist, vapor; **N.L. neut. adj. suff.** *-genum*, born from; **N.L. neut. adj.** *capeverdeinubigenum*, born of the clouds in Cape Verde

**Nomenclatural type**
Pending Genome: Curtobacterium_capeverdeinubigenum Ts

**Description**
*Curtobacterium capeverdeinubigenum* MPC1 grows as white -yellow, round colony, diameter 1 mm, was isolated from cloud water sample WW85

**Classification**
*Bacteria* » *Actinomycetota* » *Actinomycetes* » *Micrococcales* » *Microbacteriaceae* » *Curtobacterium* » *Curtobacterium capeverdeinubigenum*

**Registry URL**
https://seqco.de/i:55978

#### Species *Curtobacterium nubigenum*

**Etymology**
[nu.bi.ge'num] **L. fem. n.** *nubes*, a cloud, mist, vapor; **N.L. neut. adj. suff.** *-genum*, born from; **N.L. neut. adj.** *nubigenum*, born from the clouds

**Nomenclatural type**
INSDC Nucleotide: JBPGPG000000000 Ts

**Description**
The *Curtobacterium nubigenum* MPC39 strain has round, yellow, 2 mm diameter colonies, it was isolated from cloud water sample WW29

**Classification**
*Bacteria* » *Actinomycetota* » *Actinomycetes* » *Micrococcales* » *Microbacteriaceae* » *Curtobacterium* » *Curtobacterium nubigenum*

**Registry URL**
https://seqco.de/i:55980

#### Species *Deinococcus monteverdensis*

**Etymology**
[mon.te.ver.den'sis] **N.L. masc. adj.** *monteverdensis*, of Monte Verde, on Cape Verde Islands

**Nomenclatural type**
Pending Genome: SAMN48999379 Ts

**Description**
This MAG was obtained from sequening the cloud water sample WW85 collected on Mount Verde on the Cape Verde Islands

**Classification**
*Bacteria* » *Deinococcota* » *Deinococci* » *Deinococcales* » *Deinococcaceae* » *Deinococcus* » *Deinococcus monteverdensis*

**Registry URL**
https://seqco.de/i:55985

#### Species *Deinococcus nubigenus*

**Etymology**
[nu.bi.ge'nus] **L. fem. n.** *nubes*, a cloud, mist, vapor; **N.L. masc. adj. suff.** *-genus*, born from; **N.L. masc. adj.** *nubigenus*, born from the clouds

**Nomenclatural type**
Pending Genome: Deinococcus_nubigenus Ts

**Description**
The *Deinococcus nubigenus* MPC36 strain has round red colonies, 1 mm in diameter, was isolated from cloud water sample WW29

**Classification**
*Bacteria* » *Deinococcota* » *Deinococci* » *Deinococcales* » *Deinococcaceae* » *Deinococcus* » *Deinococcus nubigenus*

**Registry URL**
https://seqco.de/i:55979

#### Species *Duganella nubigena*

**Etymology**
[nu.bi.ge'na] **L. fem. n.** *nubes*, a cloud, mist, vapor; **N.L. fem. adj. suff.** *-gena*, born from; **N.L. fem. adj.** *nubigena*, born from the clouds

**Nomenclatural type**
Pending Genome: SAMN48999364 Ts

**Description**
This MAG was obtained from sequening the cloud water sample WW29 collected on Mount Verde on the Cape Verde Islands

**Classification**
*Bacteria* » *Pseudomonadota* » *Betaproteobacteria* » *Burkholderiales* » *Oxalobacteraceae* » *Duganella* » *Duganella nubigena*

**Registry URL**
https://seqco.de/i:55983

#### Species *Methylobacterium nubigenum*

**Etymology**
[nu.bi.ge'num] **L. fem. n.** *nubes*, a cloud, mist, vapor; **N.L. neut. adj. suff.** *-genum*, born from; **N.L. neut. adj.** *nubigenum*, born from the clouds

**Nomenclatural type**
Pending Genome: SAMN48999391 Ts

**Description**
This MAG was obtained from sequening the cloud water sample WW85 collected on Mount Verde on the Cape Verde Islands

**Classification**
*Bacteria* » *Pseudomonadota* » *Alphaproteobacteria* » *Hyphomicrobiales* » *Methylobacteriaceae* » *Methylobacterium* » *Methylobacterium nubigenum*

**Registry URL**
https://seqco.de/i:55988

#### Species *Noviherbaspirillum nubigenum*

**Etymology**
[nu.bi.ge'num] **L. fem. n.** *nubes*, a cloud, mist, vapor; **N.L. neut. adj. suff.** *-genum*, born from; **N.L. neut. adj.** *nubigenum*, born from the clouds

**Nomenclatural type**
Pending Genome: SAMN48999387 Ts

**Description**
This MAG was obtained from sequening the cloud water sample WW85 collected on Mount Verde on the Cape Verde Islands

**Classification**
*Bacteria* » *Pseudomonadota* » *Betaproteobacteria* » *Burkholderiales* » *Oxalobacteraceae* » *Noviherbaspirillum* » *Noviherbaspirillum nubigenum*

**Registry URL**
https://seqco.de/i:55987

#### Species *Novosphingobium monteverdense*

**Etymology**
[mon.te.ver.den'se] **N.L. neut. adj.** *monteverdense*, of Monte Verde, on Cape Verde Islands

**Nomenclatural type**
Pending Genome: SAMN48999352 Ts

**Description**
This MAG was obtained from sequencing the blank water BW13 used to clean between cloud water samplings

**Classification**
*Bacteria* » *Pseudomonadota* » *Alphaproteobacteria* » *Sphingomonadales* » *Erythrobacteraceae* » *Novosphingobium* » *Novosphingobium monteverdense*

**Registry URL**
https://seqco.de/i:55982

#### Species *Pantoea nubigena*

**Etymology**
[nu.bi.ge'na] **L. fem. n.** *nubes*, a cloud, mist, vapor; **N.L. fem. adj. suff.** *-gena*, born from; **N.L. fem. adj.** *nubigena*, born from the clouds

**Nomenclatural type**
Pending Genome: SAMN48999403 Ts

**Description**
This MAG was obtained from sequening the cloud water sample WW8 collected on Mount Verde on the Cape Verde Islands

**Classification**
*Bacteria* » *Pseudomonadota* » *Gammaproteobacteria* » *Enterobacterales* » *Erwiniaceae* » *Pantoea* » *Pantoea nubigena*

**Registry URL**
https://seqco.de/i:55990

#### Species *Ralstonia monteverdensis*

**Etymology**
[mon.te.ver.den'sis] **N.L. fem. adj.** *monteverdensis*, of Monte Verde, on Cape Verde Islands

**Nomenclatural type**
Pending Genome: SAMN48999346 Ts

**Description**
This MAG was obtained from sequening the blank water BW13 used to clean between cloud water samplings

**Classification**
*Bacteria* » *Pseudomonadota* » *Betaproteobacteria* » *Burkholderiales* » *Burkholderiaceae* » *Ralstonia* » *Ralstonia monteverdensis*

**Registry URL**
https://seqco.de/i:55981

#### Species *Sphingomonas nubigena*

**Etymology**
[nu.bi.ge'na] **L. fem. n.** *nubes*, a cloud, mist, vapor; **N.L. fem. adj. suff.** *-gena*, born from; **N.L. fem. adj.** *nubigena*, born from the clouds

**Nomenclatural type**
Pending Genome: SAMN48999366 [Ts]

**Description**
This MAG was obtained from sequening the cloud water sample WW29 collected on Mount Verde on the Cape Verde Islands

**Classification**
*Bacteria* » *Pseudomonadota* » *Alphaproteobacteria* » *Sphingomonadales* » *Sphingomonadaceae* » *Sphingomonas* » *Sphingomonas nubigena*

**Registry URL**
https://seqco.de/i:55984

#### Species *Spirosoma nubigenum*

**Etymology**
[nu.bi.ge'num] **L. fem. n.** *nubes*, a cloud, mist, vapor; **N.L. neut. adj. suff.** *-genum*, born from; **N.L. neut. adj.** *nubigenum*, born from the clouds

**Nomenclatural type**
Pending Genome: SAMN48999396 [Ts]

**Description**
This MAG was obtained from sequening the cloud water sample WW85 collected on Mount Verde on the Cape Verde Islands

**Classification**
*Bacteria* » *Bacteroidota* » *Cytophagia* » *Cytophagales* » *Spirosomataceae* » *Spirosoma* » *Spirosoma nubigenum*

**Registry URL**
https://seqco.de/i:55989

#### Species *Variovorax nubigenus*

**Etymology**
[nu.bi.ge'nus] **L. fem. n.** *nubes*, a cloud, mist, vapor; **N.L. masc. adj. suff.** *-genus*, born from; **N.L. masc. adj.** *nubigenus*, born from the clouds

**Nomenclatural type**
Pending Genome: SAMN48999386 [Ts]

**Description**
This MAG was obtained from sequening the cloud water sample WW85 collected on Mount Verde on the Cape Verde Islands

**Classification**
*Bacteria* » *Pseudomonadota* » *Betaproteobacteria* » *Burkholderiales* » *Comamonadaceae* » *Variovorax* » *Variovorax nubigenus*

**Registry URL**
https://seqco.de/i:55986
